# Activity-based cell sorting reveals resistance of functionally degenerate *Nitrospira* during a press disturbance in nitrifying activated sludge

**DOI:** 10.1101/2021.04.24.441178

**Authors:** Maxwell B.W. Madill, Yaqian Luo, Pranav Sampara, Ryan M. Ziels

**Author notes:** Contributed equally to this work (Maxwell B.W. Madill, Yaqian Luo).

## Abstract

Managing and engineering activated sludge wastewater treatment microbiomes for low-energy nitrogen removal requires process control strategies to stop the oxidation of ammonium at nitrite. Our ability to out-select nitrite oxidizing bacteria (NOB) from activated sludge is challenged by their metabolic and physiological diversity, warranting measurements of their *in situ* physiology and activity under selective growth pressures. Here, we examined the stability of nitrite oxidation in activated sludge during a press disturbance induced by treating a portion of return activated sludge with a sidestream flow containing free ammonia (FA) at 200 mg NH_3_-N/L. The nitrite accumulation ratio peaked at 42% by day 40 in the experimental bioreactor with the press disturbance, while it did not increase in the control bioreactor. A subsequent decrease in nitrite accumulation within the experimental bioreactor coincided with shifts in dominant *Nitrospira* 16S rRNA amplicon sequence variants (ASVs). We applied bioorthogonal non-canonical amino acid tagging (BONCAT) coupled with fluorescence activated cell sorting (FACS) to investigate changes in the translational activity of NOB populations throughout batch exposure to FA. BONCAT-FACS confirmed that the single *Nitrospira* ASV washed-out of the experimental bioreactor had reduced translational activity following exposure to FA, whereas the two *Nitrospira* ASVs that emerged after process acclimation were not impacted by FA. Thus, the coexistence of functionally degenerate and physiologically resistant *Nitrospira* populations provided resilience to the nitrite-oxidizing function during the press disturbance. These results highlight how BONCAT-FACS can resolve ecological niche differentiations within activated sludge and inform strategies to engineer and control microbiome function.

**Importance:** Nitrogen removal from activated sludge wastewater treatment systems is an energy-intensive process due to the large aeration requirement for nitrification. This energy footprint could be minimized with engineering control strategies that wash-out nitrite oxidizing bacteria (NOB) to limit oxygen demands. However, NOB populations can have a high degree of physiological diversity, and it is currently difficult to decipher the behavior of individual taxa during applied selective pressures. Here, we utilized a new substrate analog probing approach to measure the activity of NOB at the cellular translational level in the face of an applied press disturbance to the activated sludge process. Substrate analog probing corroborated the time-series reactor sampling, showing that coexisting and functionally redundant *Nitrospira* provided resilience to the nitrite oxidation process. Taken together, these results highlight how substrate analog approaches can illuminate *in situ* ecophysiologies within shared niches, and can inform strategies to improve microbiome engineering and management.

## Introduction

Relating the in situ physiological responses of individual taxa in the face of environmental perturbations to the resulting microbial community structure and function remains a critical challenge for controlling and engineering microbiomes for desirable ecological processes and outcomes (1–3). Biological wastewater treatment processes are ideal ecosystems to explore such relationships, as environmental conditions can be manipulated, and the community function monitored, in a relatively controlled manner (4). A major goal in the wastewater industry is to engineer microbial bioprocesses to achieve energy-efficient, or even net energy-positive, wastewater treatment (5). A foundational component of achieving that goal is the optimization of mainstream biological nitrogen removal processes, as conventionally this process is the largest consumer of energy and exogenous organic carbon within wastewater treatment plants (WWTPs) (6, 7). Realizing energy-efficient nitrogen removal requires highly finessed and sustained modulation of abundances, activities and interactions of key microbial functional groups to effectively control the global community function and engage the desired nitrogen removal pathway(s) (8, 9). As such pathways typically impose inherent energetic and/or metabolic constraints (10, 11), and often challenge existing community interactions (8, 9), it is critical to fully illuminate the ecophysiological diversity and mechanisms driving niche partitioning within those microbial functional groups, as well as their responses to the applied process control strategies (9).

An appealing strategy to achieve energy-efficient biological nitrogen removal is to limit the nitrification process to nitritation (i.e. oxidation of ammonium to nitrite), as the resulting nitrite can be denitrified directly (25% and 40% net energy and carbon reductions, respectively), and/or provided to anammox bacteria as a growth substrate for autotrophic nitrogen removal (60% and 100% net energy and carbon reductions, respectively) (8, 12). However, achieving stable nitritation in mainstream activated sludge (AS) stands as a major challenge limiting its successful full-scale implementation (13–15). Realizing stable nitritation in mainstream AS relies on engineering control strategies that serve as press disturbances to consistently repress and wash-out nitrite oxidizing bacteria (NOB) while maintaining the activity of ammonium oxidizing bacteria (AOB) (8, 9, 13). The efficacy of a given control strategy is therefore dependent on its ability to create a disturbance that elicits different physiological responses between AOB and NOB. Preliminary success in washing-out NOB from mainstream AS has been achieved using press disturbances that provide a high ammonium residual (13) or control the availability of dissolved oxygen (13, 16, 17) to favour the growth kinetics of AOB over NOB. Additionally, several recently-proposed control strategies have utilized the higher innate sensitivity of NOB to free-ammonia (FA) and free-nitrous acid (FNA) compared to AOB (18–20) to achieve effective NOB inhibition (21–23). Wang et al. (21) demonstrated that a press disturbance induced by exposing a fraction of return sludge to FA-rich sidestream wastewater (210 mg NH_3_-N/L) supported successful NOB wash-out in mainstream AS, with nitrite accumulation ratios reaching 80-90%. Despite its potential efficacy for supporting mainstream nitritation, there have been a limited number of studies evaluating the role of niche differentiation and physiological diversity on the stability of nitrite oxidation in the face of a press disturbance from routine FA exposure.

NOB communities in wastewater treatment often display functional degeneracy, wherein the nitrite oxidation process is distributed among several phylogenetically diverse taxa with varying auxiliary metabolic potentials (24–29). Inherent differences in nitrite oxidation biochemistry and cell morphology play core roles in supporting ecophysiological diversity between NOB genera by influencing their substrate affinities for oxygen and nitrite, and their nitrite oxidation kinetics (24, 30, 31). *Nitrospira,* a predominant NOB genus in many WWTP microbiomes (32–34), has demonstrated an extraordinary degree of functional degeneracy, with reports of highly complex and stable communities containing as many as 120 closely-related coexisting strains (26, 27, 34). Considerable ecophysiological diversity may thus exist between *Nitrospira* species/strains to support niche partitioning, which could be supported by distinct oxygen and nitrite preferences (34–36), auxiliary metabolic potentials for utilizing alternative electron acceptors and/or donors (24, 27, 34, 37, 38), and tolerances to challenging environmental conditions including FA (27, 28, 39, 40). Exhibited at both the genus and strain-level, such functional degeneracy may enable NOB communities to resist the selective pressures imparted by engineering process control strategies by recruiting functionally redundant, yet physiologically diverse, NOB members (41–43). *In situ* assessments of the metabolically active fraction of nitrifying communities are therefore critical to evaluate the efficacy of mainstream nitritation control strategies and elucidate their associated impacts on functionally degenerate NOB.

Next-generation substrate analogue probing approaches have recently emerged as powerful tools to decipher the *in situ* physiology of active cells based on their uptake of synthetic analogues of natural biomolecules (44). Bioorthogonal non-canonical amino acid tagging (BONCAT) is a nascent SAP approach to study the physiology of active cells in complex environmental microbiomes (45–48). BONCAT relies on the *in vivo* uptake and incorporation of synthetic amino acids, such as the alkyne-containing analogue of methionine, homopropargylglycine (HPG), into newly synthesized proteins via the native translational machinery, and thereby selectively labels the proteomes of translationally active cells (44, 47). HPG-labeled cells can subsequently be identified by tagging their proteins with azide-modified fluorescent dyes via azide-alkyne clickchemistry, enabling their selective recovery using fluorescence activated cell sorting (FACS) (45, 46, 48). To our knowledge, BONCAT, or its paired approach with FACS (BONCAT-FACS), have yet to be applied to study the active fractions of AS microbiomes central to wastewater treatment bioprocesses.

The objective of this study was to assess the stability of nitrite oxidation in the face of a press disturbance induced by routine exposure of recycled activated sludge with FA as an engineering control strategy to wash-out NOB. We hypothesized that certain members of the active NOB microbial community could acclimate to the applied press disturbance. Two parallel experimental and control AS sequencing batch reactors (SBRs) were operated for ~100 days to investigate the impacts of routine FA exposure as a press disturbance on the NOB community. We applied time-series 16S rRNA gene amplicon sequencing in addition to BONCAT-FACS based activity measurements to elucidate changes in the structure and *in situ* activity of the AS microbiome and nitrifying communities.

## Results

### Partial nitritation performance of activated sludge bioreactors

Two AS SBRs treated synthetic mainstream municipal wastewater over two operational phases: the Start-up Phase and the Treatment Phase (Figure S1). Periodic steady-state conditions were presumed over the last 30 days of the 270-day Start-up Phase, during which there were no significant differences in daily effluent concentrations of NH_4_^+^-N, NO_3_^-^-N, and NO_2_^-^-N between the two SBRs (*p* > 0.05), which averaged 0.1 ±0.3 mg NH_4_^+^-N/L, 19±2 mg NO_3_^-^-N/L and 0.0±0.01 mg NO_2_^-^-N/L, respectively (Figure 1). There were also no significant differences in the TSS and VSS of mixed liquor between the two systems during the Start-up Phase (*p > 0.05*; Figure S2). Thus, similar stable performance with full nitrification was achieved in both SBRs during the initial Start-up Phase, indicating effective duplication of operating conditions.

The Treatment Phase was then commenced on day 0 to assess the impacts of a press disturbance induced by routine FA-exposure of return sludge in a side-stream reactor on the nitrifying community structure and activity. Approximately 20% of the return sludge in the treatment SBR was exposed to 200 mg NH_3_-N/L as FA at a pH of 9.0 for 24 hrs before being reintroduced into the mainstream SBR on a daily basis, while the same conditions were emulated for the control SBR but without FA added to the side-stream. Additionally, ammonium nitrogen was added (160 mg NH_4_^+^-N/d) to the control SBR on a daily basis along with the return sludge to achieve equivalent nitrogen loads between the two SBRs. After 10 days of the applied press disturbance, NO_2_^-^-N began to increase in effluent of the treatment SBR, but stayed below detection in the control SBR for the remainder of the Treatment Phase (Figure 1). By day 40, effluent NO_2_^-^-N reached its peak level of 11 mg NO_2_^-^-N/L in the treatment SBR. At the same time, NH✓-N accumulated to 3.7 mg NH4-N/L in the treatment SBR between days 37 and 44, yet stayed below 1.2 mg NH4-N/L in the control SBR over the entire Treatment Phase (Figure 1). The accumulation of NO_2_^-^-N in the treatment SBR was transient, however, as the effluent concentration decreased after day 40 and reached a value below detection by day 74 (Figure 1).

**Figure 1:**
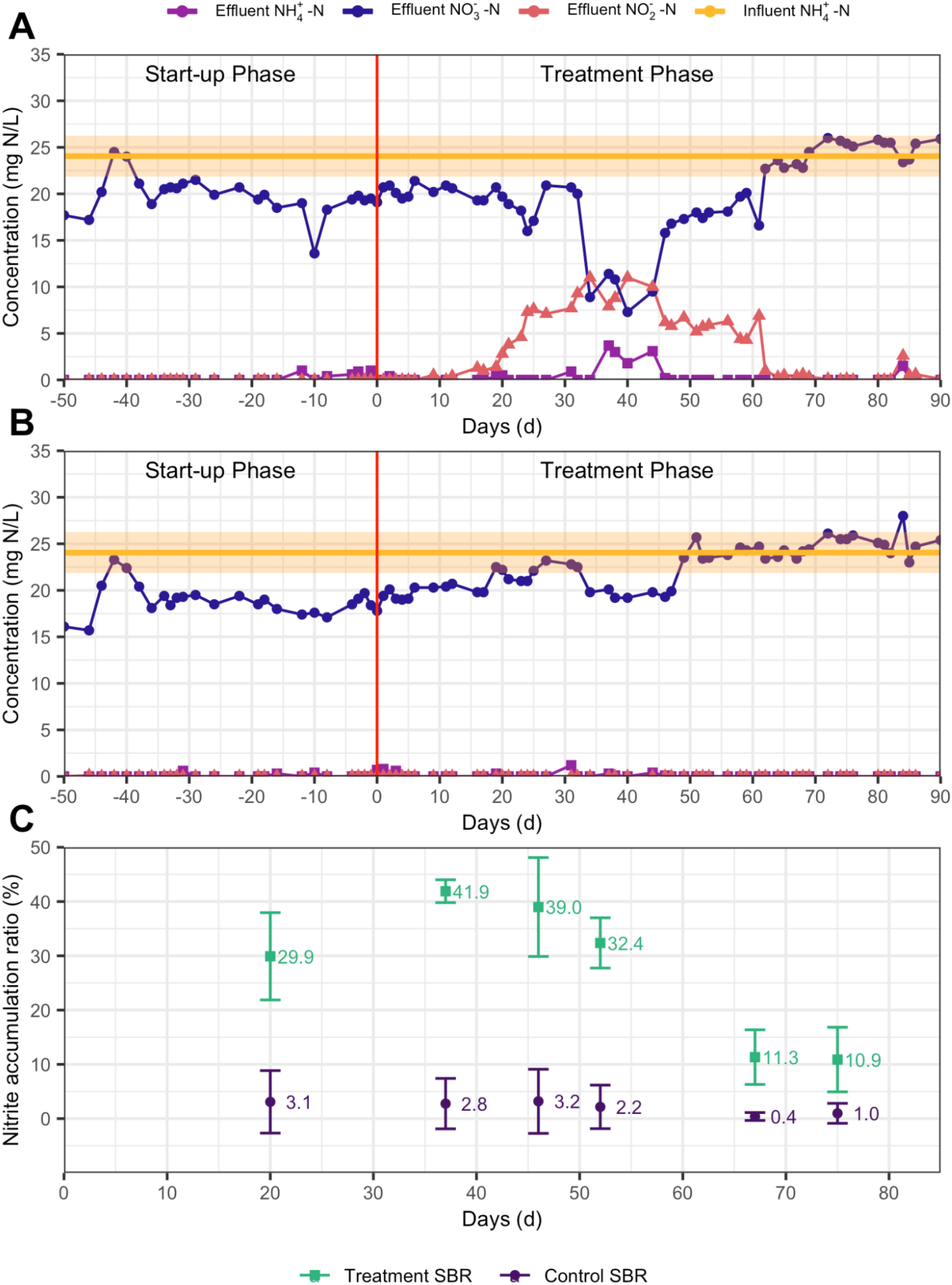
Influent and effluent nitrogen species over the two experimental phases in the (A) treatment sequencing batch reactor (SBR), and (B) control SBR. Shaded orange space represents standard deviation of influent NH_4_^+^-N. (C) Nitrite accumulation ratio (NAR, %) of treatment and control SBRs over time, based on 24 h monitoring of SBR effluents. NAR is defined as the effluent nitrite (mg N/L) divided by the effluent nitrite plus nitrate (mg N/L), and indicates the level of NOB activity suppression.

As effluent was sampled from the SBRs on a 24 h basis, and the sidestream return sludge was added once every 24 h, periodic tests were conducted to measure nitrogen species at the end of individual SBR cycles over the course of 24 h to better estimate nitrite accumulation ratios (NARs) (Figure S3). After 20 days of the press-disturbance, the 24 h average NAR was approximately 10 times higher in the treatment SBR versus the control (Figure 1C). The NAR reached its peak of 41.9% by day 37 in the treatment SBR, aligning with the observed peak in effluent NO_2_^-^-N concentrations. After day 37, the NAR decreased in the treatment SBR, reaching its lowest observed level of 10.9% on day 75. The NAR of the control SBR stayed below 3.2% in the entire Treatment Phase (Figure 1C). This suggests that the nitrite oxidation function was resilient to the press disturbance of routine FA exposure, as the extent of nitrite oxidation inhibition was not sustained after about 40 days.

### 3.2 Microbial community acclimation to routine FA-exposure

Microbial 16S rRNA gene amplicons were denoised into amplicon sequence variants (ASVs) to provide a high-resolution (49, 50) view of how routine FA-exposure impacted the community structure. A total of 3.01 million chimera-free quality-filtered merged reads (Table S1) were denoised into 6,694 ASVs. Over 95% of the 16S rRNA gene amplicons at all time points in both SBRs were comprised of the 8 phyla: *Proteobacteria, Bacteroidetes, Chloroflexi, Nitrospirae, Planctomycetes, Verrucomicrobia, Acidobacteria,* and *Cyanobacteria* (Figure S4). Between 53% to 70% percent of 16S rRNA amplicons were represented by 20 genera across all samples (Figure 2). Even at the broad genus level of resolution, there were apparent differences in community profiles between the treatment and control SBRs over time (Figure 2), indicating that routine FA exposure altered the structure of the AS microbiome.

**Figure 2:**
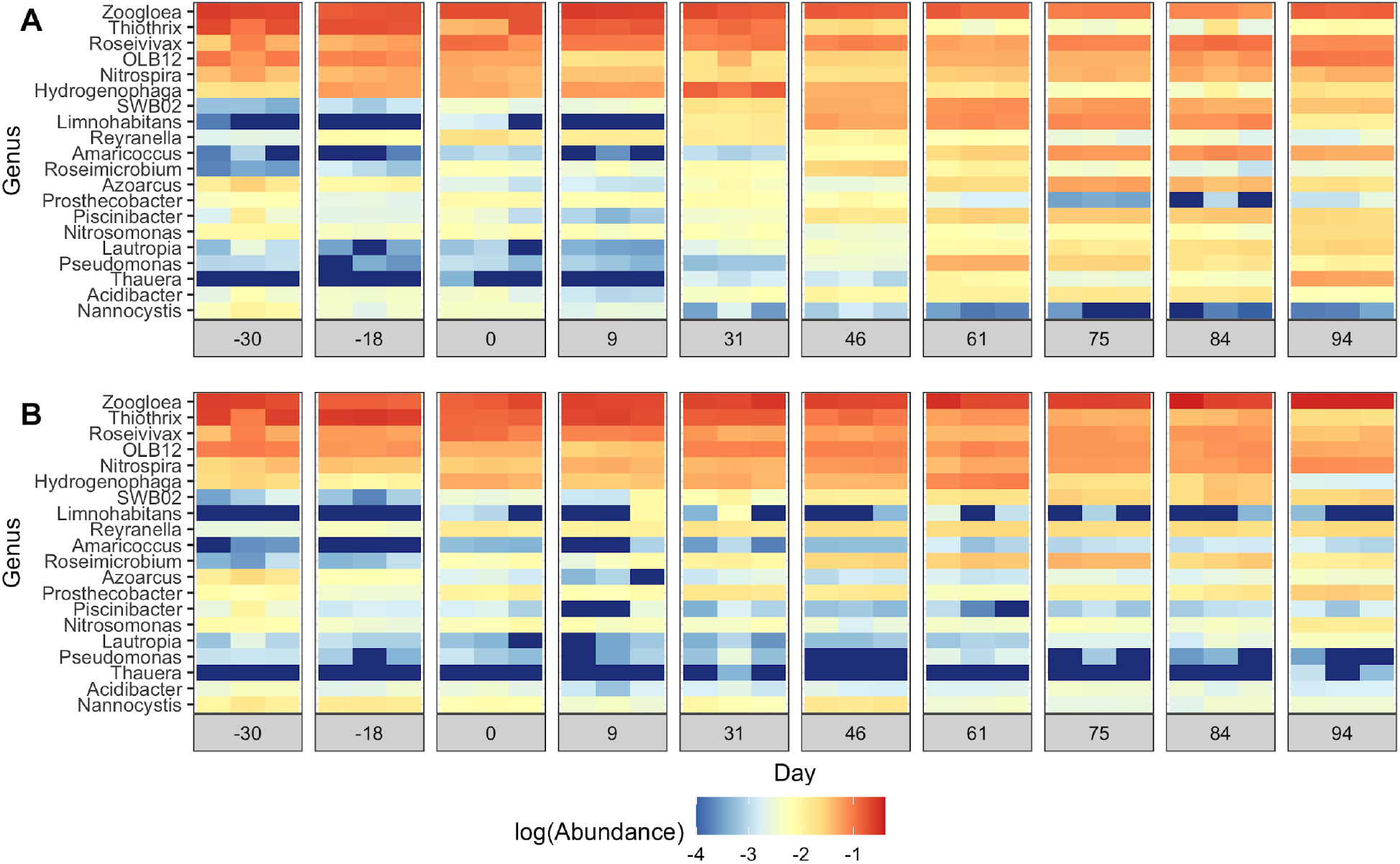
Heatmap of log-scaled relative abundance of top 20 most abundant genera in the (A) treatment SBR and (B) control SBR over the two operational phases. Sequencing results are shown for triplicate DNA extractions on each sampling date. The genera are ordered from highest to lowest cumulative abundance across samples.

At the ASV level, FA-exposure of return sludge led to significant differences in community structure between the two SBRs over time (*R*^2^ = 0.54, p < 0.001, adonis). Principal coordinates analysis (PCoA) of cumulative sum-scaled ASV read counts revealed that community profiles in the two SBRs were similar until day 9, after which the treatment SBR community diverged from the control (Figure 3). Differential abundance analysis showed that there were no statistically different ASVs between the two SBRs on day 0 (*p* > 0.01, DESeq2), indicating that they were well replicated in the Start-up Phase. By day 46 of the Treatment Phase, around the time that nitrite peaked in the treatment SBR (Figure 1), 105 ASVs spanning 55 genera were differentially abundant between the two SBRs (*p* < 0.01, DESeq2; Figure S5). The number of ASVs with significant differential abundance between the SBRs continued to increase to a maximum of 166 on day 75 of the Treatment Phase (Figure S5).

**Figure 3:**
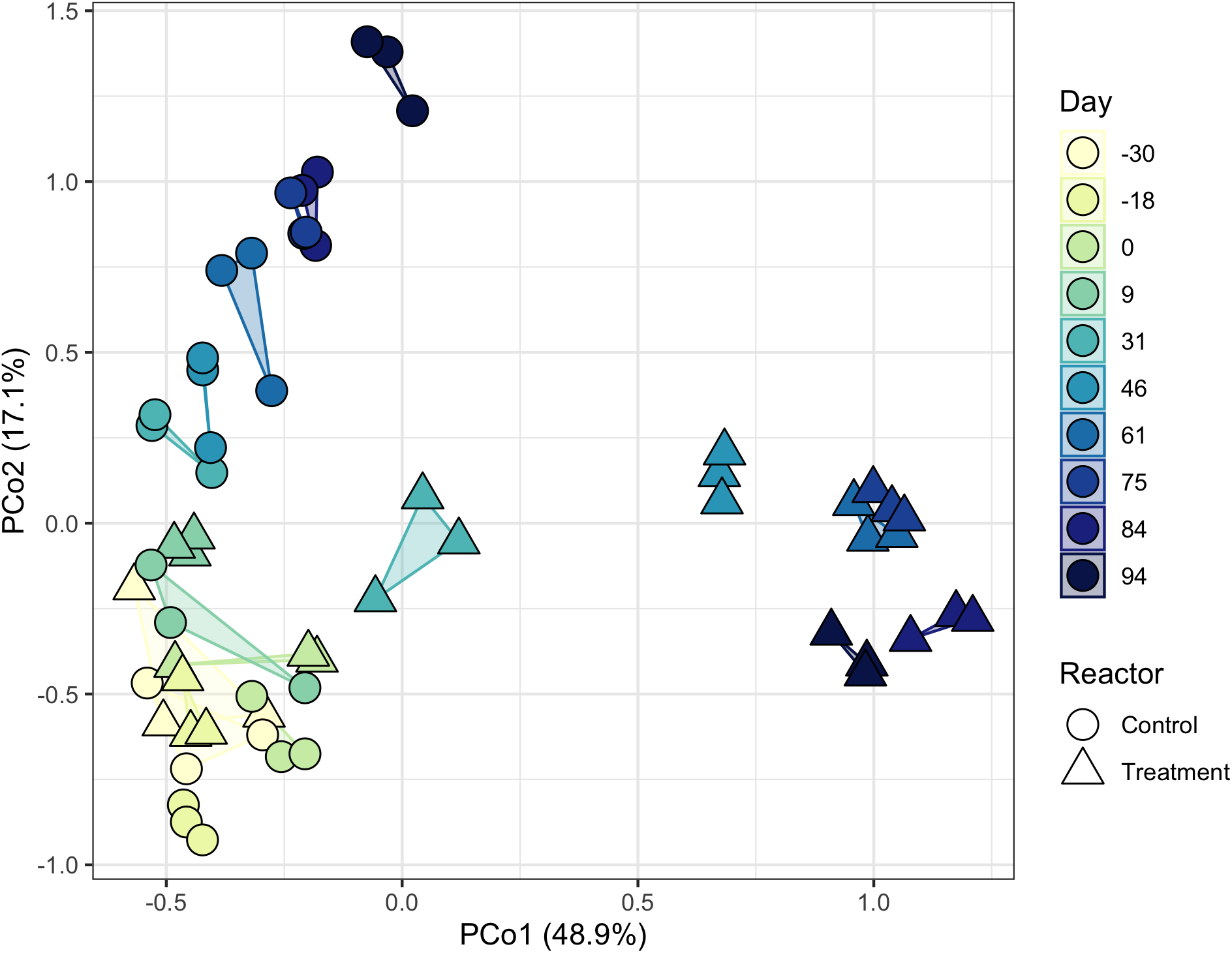
Principal coordinates analysis (PCoA) of Bray-Curtis dissimilarities between cumulative sum scaled (CSS) read counts of 16S rRNA ASVs in both SBRs over time. The marker fill represents reactor operating day, and the marker shape corresponds to the SBR. Triplicate DNA extractions for each time point are indicated by a shared polygon. The percentages in parentheses represent the fraction of variance explained by that axis.

*Nitrospira* and *Nitrosomonas* were the only putative NOB and AOB populations detected in the SBRs, respectively (Figure 2). As these reactors were fed with synthetic wastewater, it is likely that these populations originated from the inoculum. Six dominant *Nitrosomonas* ASVs were detected in both SBRs over the two experimental phases (Figure S6). Until day 84, the total *Nitrosomonas* abundance was less than 1% in both SBRs, but increased to over 1.5% in both by day 94 (Figure S12). One *Nitrosomonas* ASV (ASV_36) was differentially abundant between the two SBRs on days 75 and 94 (p < 0.01, DESeq2). Three *Nitrospira* ASVs were detected within the two SBRs (Figure 4). In particular, ASV_8 was the dominant *Nitrospira* ASV in both SBRs during the Start-up Phase (before day 0), accounting for 3.3%±0.8% of 16S rRNA genes on average (Figure 4). By day 46, ASV_8 decreased to 1.6%±0.1% in the treatment SBR, while it increased to 6.4%±0.3% within the control. This decrease in ASV_8 abundance coincided with the peak in nitrite accumulation in the treatment SBR (Figure 1a). The abundance of ASV_8 did not significantly change after day 46 for the remainder of the experiment in the treatment SBR (p > 0.01, DESeq2). In contrast, two other *Nitrospira* ASVs (ASV_32 and ASV_47; 99.7% sequence similarity to each other; 94.2% and 94.5% sequence similarities to ASV_8, respectively) were sporadically detected in both SBRs at abundances below 0.4% until day 46, and then increased to a maximum of 1.8%±0.4% and 1.2%±0.3% in the treatment SBR by day 84, respectively, but stayed below 0.3% in the control (Figure 4). The abundance of ASV_32 and ASV_47 were significantly higher in the treatment SBR relative to the control SBR by the end of the experiment, while ASV_8 was significantly lower (both days 84 and 94; *p* < 0.01, DESeq2). Phylogenetic analysis based on partial 16S rRNA gene sequences (Figure S7) revealed that ASV_8 was most closely related to *Nitrospira lenta* within lineage II, whereas ASV_32 and ASV_47 were clustered within lineage I of *Nitrospira* (51,52).

**Figure 4:**
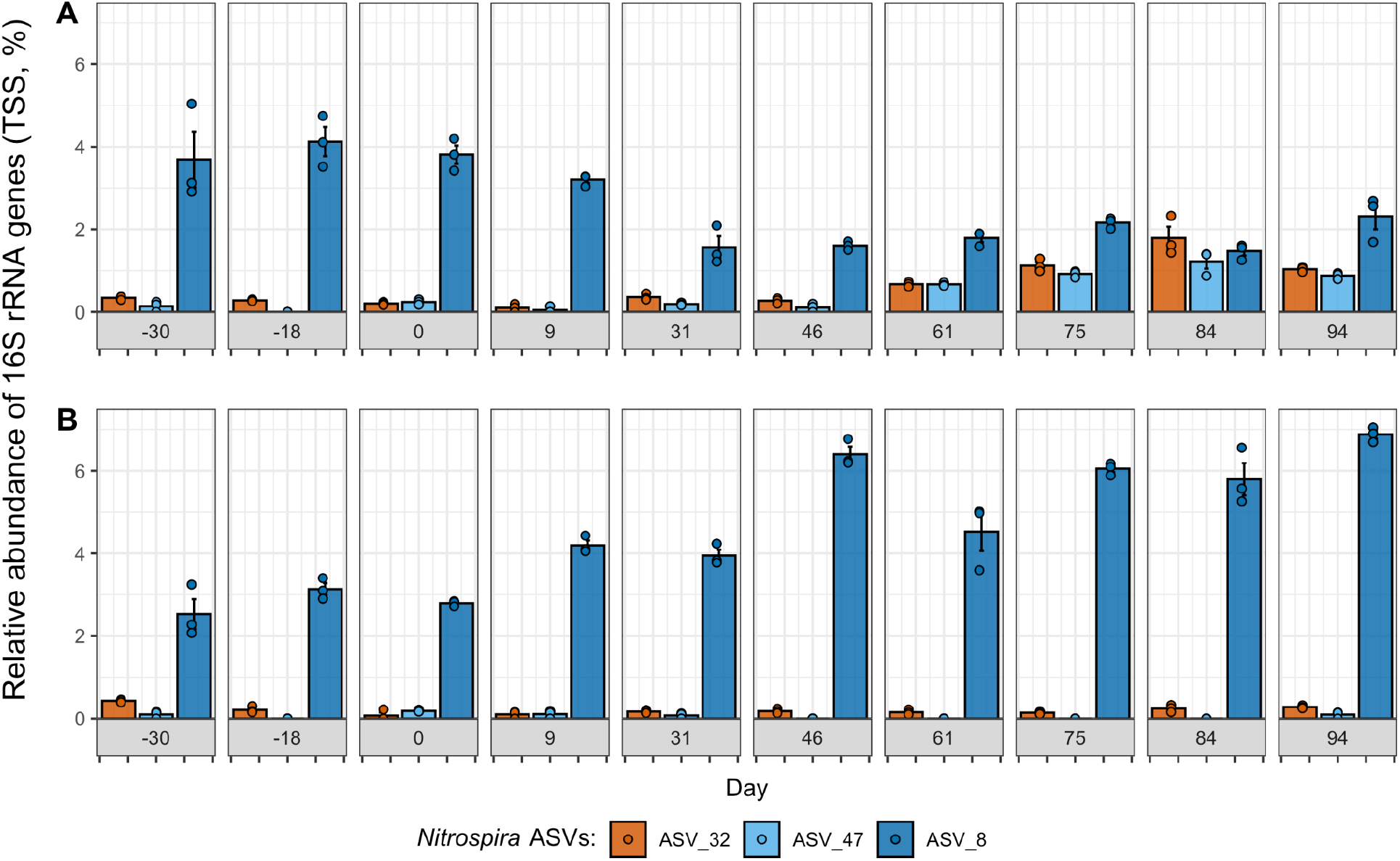
Relative abundance (total sum scaled, TSS) of three dominant *Nitrospira* ASVs over time in the (A) treatment SBR and (B) control SBR. Only *Nitrospira* ASVs detected in more than one sample are shown. Results are shown for triplicate DNA extractions on each day, with the coloured points showing the relative abundance of each ASV in each DNA extraction, the coloured bar showing the mean relative abundance, and the error bars showing the standard error of the mean.

### 3.3 Substrate analog probing of active nitrifying populations

To decipher the impact of the applied press disturbance on the activity and *in situ* physiology of nitrifying populations within mainstream AS, we conducted BONCAT-FACS on samples collected from nitrifying microcosms seeded with SBR mixed liquor preceding return sludge treatment (R), as well as return sludge from the beginning (15 min after start; S1) and end (24 h after start; S2) of the side-stream return sludge treatment processes for each SBR (Figure 5). Preliminary validation microcosms confirmed the sensitivity of BONCAT in labeling only active cells that had incorporated HPG (Figure S8). Comparison of FACS data between the two SBRs (Figures S9 and S10) revealed that translationally active (i.e. BONCAT+) cell fractions were significantly lower in the S1 and S2 nitrifying microcosms seeded from the FA-exposed treatment SBR relative to those of the control SBR (*p* < 0.05, t-test), but that no significant difference in BONCAT+ cell fractions was observed in the R microcosms seeded with mixed-liquor preceding sludge treatment. (Figure 5). Within the treatment SBR nitrifying microcosms, the fraction of BONCAT+ cells in the S1 microcosm was 25% lower than that of the R microcosm without FA-exposure, but the this was not significant (*p* = 0.078; Figure 5). No further reduction in the fraction of BONCAT+ cells was observed between microcosms seeded with return sludge from the beginning (S1) and end (S2) of the FA-exposure process (Figure 5). In contrast, the BONCAT+ cell fraction in the microcosm seeded with control SBR return sludge after 24 hrs of its sludge treatment (S2) was significantly higher by 63% compared to that at the beginning of its sludge treatment process (S1; *p*=0.045; Figure 5).

**Figure 5:**
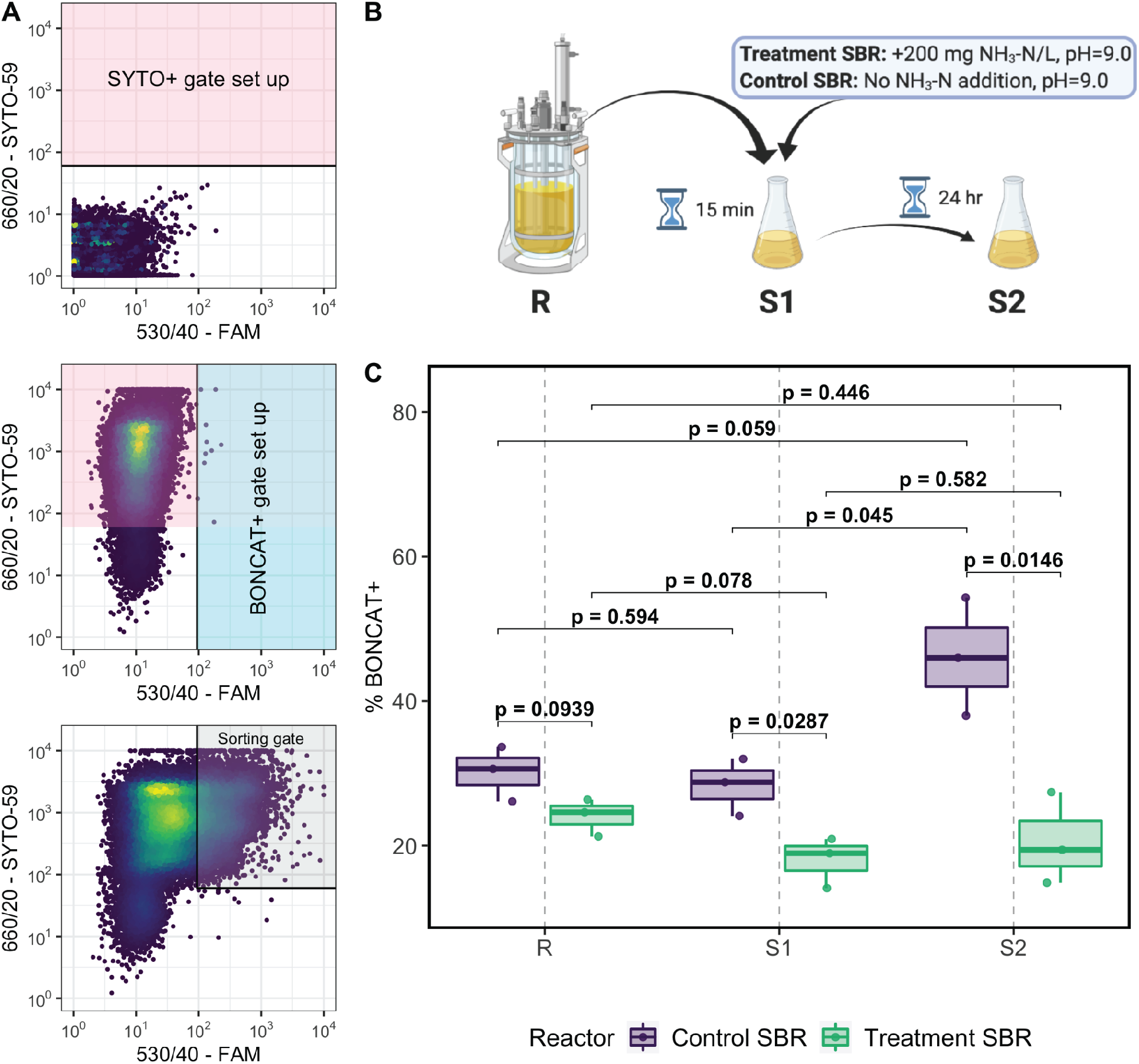
Application of BONCAT-FACS to aerobic nitrifying microcosm samples. (A) Sorting gates were established based on the detection of SYTO+ cells from background particles using unstained HPG-negative cells (top panel), and BONCAT+ cells from non-HPG labelled cells based on their FAM picolyl azide dye fluorescence using SYTO59 stained HPG-negative cells (middle panel), allowing less than 0.5% false positives in each gate. BONCAT+ cell fractions in each HPG-incubated sample were calculated as the fraction of SYTO+ cells residing in the sorting gate (bottom panel). (B) Overview of the return sludge treatment processes showing the origin of samples used to seed the nitrifying microcosms. (C) Comparison of BONCAT+ cell fractions detected in samples prepared from the nitrifying microcosms seeded with mixed-liquor preceding return sludge treatment (R), return sludge after 15 min of side-stream treatment (S1), and return sludge after 24 h of side-stream treatment (S2) from the treatment SBR (bottom) and control SBR (top). Black brackets are shown for comparisons of BONCAT+ cell fractions made between samples, with *p* values calculated using independent t-tests.

The microbial community composition observed in the BONCAT+ cell fractions could be attributed to changes in cellular translational activity, as well as any changes that occurred in the bulk community composition throughout the incubation and/or sample processing steps upstream of FACS. To delineate these impacts, we conducted 16S rRNA gene amplicon sequencing on triplicate bulk (i.e. pre-homogenized), post-homogenized and click-chemistry labelled samples from each microcosm, in addition to pre-homogenized bulk samples from HPG-negative control microcosms and SBR mixed liquors prepared with different DNA extraction and PCR amplification procedures. The low-biomass DNA extraction method (prepGEM kit) and the two-step 16S rRNA gene PCR amplification, which were both used to prepare the BONCAT+ libraries, both showed impacts on the community composition relative to respective samples prepared for time-series analysis (i.e. FastDNA Soil kit with one-step PCR; Figure S11). For this reason, the community compositions measured in the BONCAT+ samples were not compared to time-series samples with different DNA extraction and amplification procedures. PCoA revealed that bulk samples from microcosms incubated without HPG were similar to those with HPG, indicating that HPG did not alter the community structure during the 3 h incubation (Figure 6). Bulk samples from the control SBR microcosms (R, S1 and S2) were generally clustered with the mixed liquor sampled directly from the SBR (Figure 6B). In contrast, bulk samples from the treatment SBR microcosms diverged slightly from the bulk SBR community after 24 hrs of FA-exposure (e.g. S2 microcosms; Figure 6A), suggesting that the community composition was altered by exposure to FA. The community compositions of the post-homogenized samples were not identical to the bulk microcosm samples, which could be attributed to cell lysis or the removal of extracellular DNA during the homogenization and click-labeling procedures (45) (Figure 6). Due to the aforementioned findings, the BONCAT+ cell fractions were only compared to their corresponding post-homogenized microcosm samples processed with the same low-biomass DNA extraction method.

**Figure 6.**
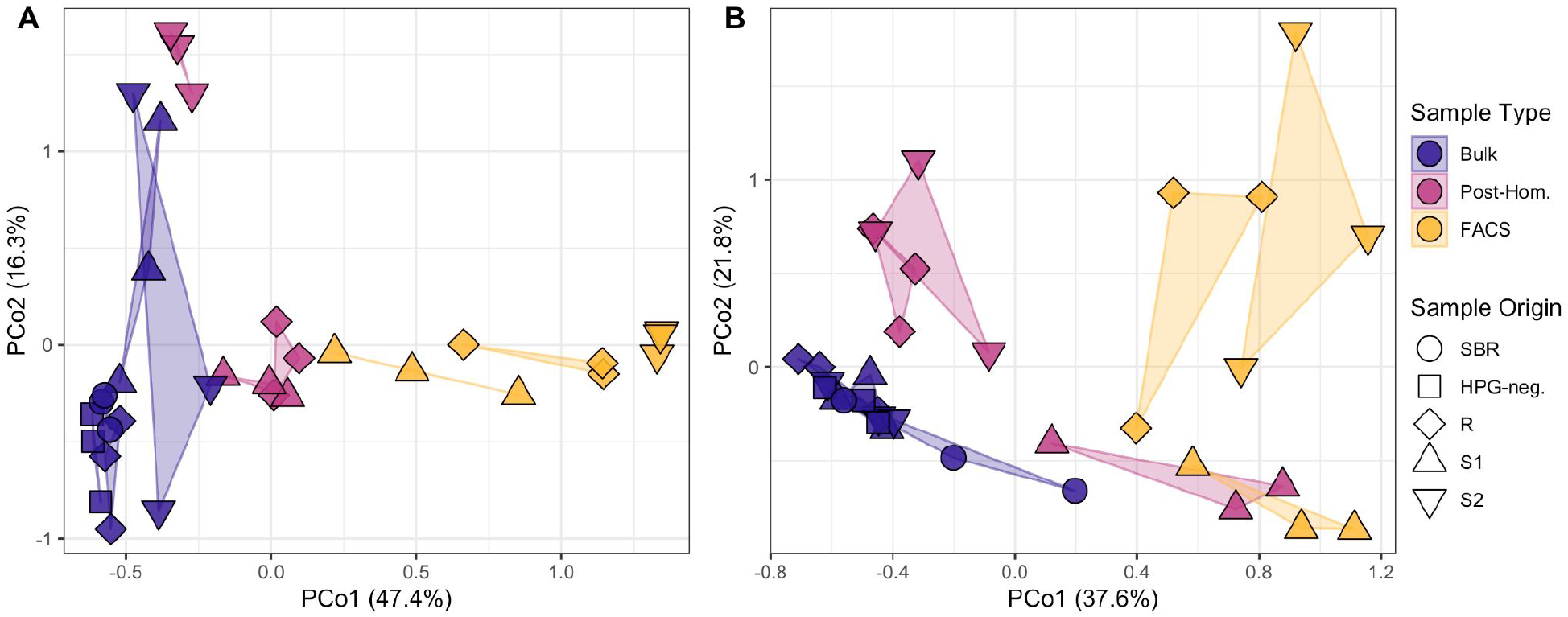
Principal coordinates analysis (PCoA) of Bray-Curtis dissimilarities between cumulative sum scaled (CSS) read counts of 16S rRNA ASVs measured in Bulk, Post-Homogenized (i.e. Post-Hom.) and BONCAT+ (i.e. FACS) samples prepared from nitrifying microcosms seeded with mixed liquor preceding side stream treatment (R), mixed liquor after 15 min of side stream treatment (S1), and mixed liquor after 24 h of side stream treatment (S2), from (A) the treatment SBR and (B) the control SBR. Bulk samples from the HPG-negative R control microcosms (i.e. HPG-neg.) and bulk mixed liquor samples directly from the SBRs (i.e. SBR) were also included. The marker fill represents the preparation procedure for each sample and the marker shape represents the sample origin (microcosm type or SBR mixed liquor). Triplicate samples are indicated by a shared polygon. The percentages in parentheses represent the fraction of variance explained by that coordinate axis.

Comparing the abundance of taxa in a BONCAT+ cell fraction to that in its corresponding bulk community prior to FACS can identify changes in translational activity at the population level (45, 48). For both the treatment and control SBRs, PCoA showed that the largest distance between post-homogenized community compositions and BONCAT+ cell fractions was observed in microcosms seeded with S2 biomass after 24 hrs of FA-exposure (Figure 6). Differential abundance analysis identified 8 and 0 differentially abundant ASVs (*p* < 0.01, DESeq2) in the BONCAT+ fractions of the treatment and control SBR mixed-liquor seeded (R) microcosms, respectively, and 0 differentially abundant ASVs in the BONCAT+ fractions of both SBR microcosms seeded with S1 biomass from the start of return sludge treatment. Conversely, for microcosms seeded with S2 biomass after 24 h of return sludge treatment, 56 and 26 ASVs were differentially abundant in the BONCAT+ fractions of the treatment SBR and control SBR samples, respectively. Within the S2 microcosm BONCAT+ fraction from the treatment SBR, the 56 differentially abundant ASVs spanned 26 genera, where 34% of the ASVs were significantly enriched and 66% were significantly reduced, relative to the post-homogenized community (Figure 7). In contrast, the 26 differentially abundant ASVs identified in the S2 microcosm BONCAT+ fraction from the control SBR, which spanned 13 genera, were all (100%) enriched relative to the post-homogenized community (Figure 7). These results suggest that side-stream return sludge treatment caused distinct shifts in the translationally active fraction of the communities in both SBRs, with FA-exposure negatively impacting the activity of a greater number of taxa compared to a similar incubation without FA.

**Figure 7:**
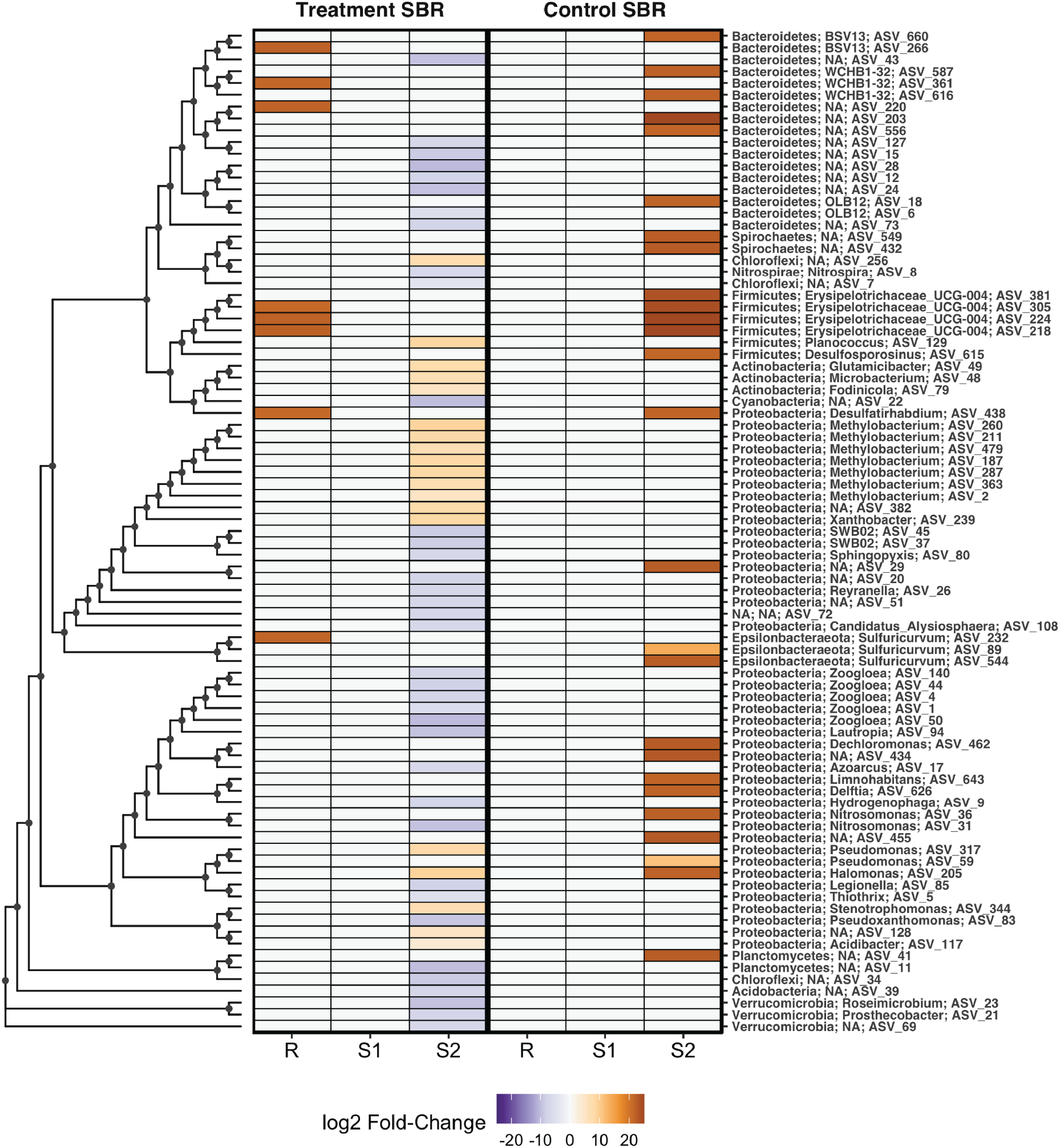
Log_2_ fold-change in abundance of all differentially abundant ASVs detected in comparisons between BONCAT+ (i.e. FACS) and Post-Homogenized (i.e. Post-Hom.) fractions prepared from nitrifying microcosms seeded with mixed-liquor preceding return sludge-treatment (R), return sludge after 15 min of side-stream treatment (S1), and return sludge after 24 h of side-stream treatment (S2) from the treatment SBR (left panel) and control SBR (right panel). For each microcosm, log_2_ fold-changes are shown only for differentially abundant ASVs, as determined by comparing ASV abundance in triplicate DNA extracts of each fraction using DESeq2 with an adjusted significance level of *p* < 0.01. Log_2_ fold-changes that were not significant are set to ‘0’ for visualization purposes. ASVs are ordered based on their phylogenetic distances estimated through multiple sequence alignment using the DECIPHER package (v.2.14.0) (53), and the phylogenetic tree was constructed using a maximum likelihood approach in the phangorn package (v.2.5.5) (54) in the R environment. Tree tip labels (right side of the heatmap) denote the phylum and genus level classifications of each ASV, where NA denotes an unknown taxonomic identity at that level.

As the BONCAT microcosms were amended with NH_4_^+^-N as the sole electron donor, it was possible to assess the impact of return sludge treatment on the translational activity of nitrifying populations. The three dominant *Nitrospira* ASVs and the top two dominant *Nitrosomonas* ASVs detected in both SBR BONCAT microcosm sample sets corresponded to the same dominant ASVs detected within both SBR mixed liquors on day 94 of the time-series sampling, when the microcosms were established (Figure 8, Figure S12). The two dominant *Nitrosomonas* ASVs (ASV_31 and ASV_36) were both differentially abundant in the BONCAT+ fractions of S2 microcosms (*p* < 0.01, DESeq2; Figure S12), while the observed differences varied by SBR. *Nitrosomonas* ASV_31 was significantly reduced in the BONCAT+ fraction of the S2 microcosm from the treatment SBR, while *Nitrosomonas* ASV_36 was significantly enriched in the BONCAT+ fraction of the S2 microcosm of the control SBR (Figure S13). Similar to *Nitrosomonas,* the only significant differential abundance in BONCAT+ fractions for *Nitrospira* ASVs occurred in a S2 microcosm (Figure 8). *Nitrospira* ASV_8 was the only differentially abundant NOB in BONCAT+ fractions of both SBR microcosms, and was significantly reduced in the S2 microcosm of the treatment SBR (*p* < 0.01, DESeq2). These results indicate that significant reductions in translational activity of nitrifying populations were only observed in return sludge exposed to FA.

**Figure 8:**
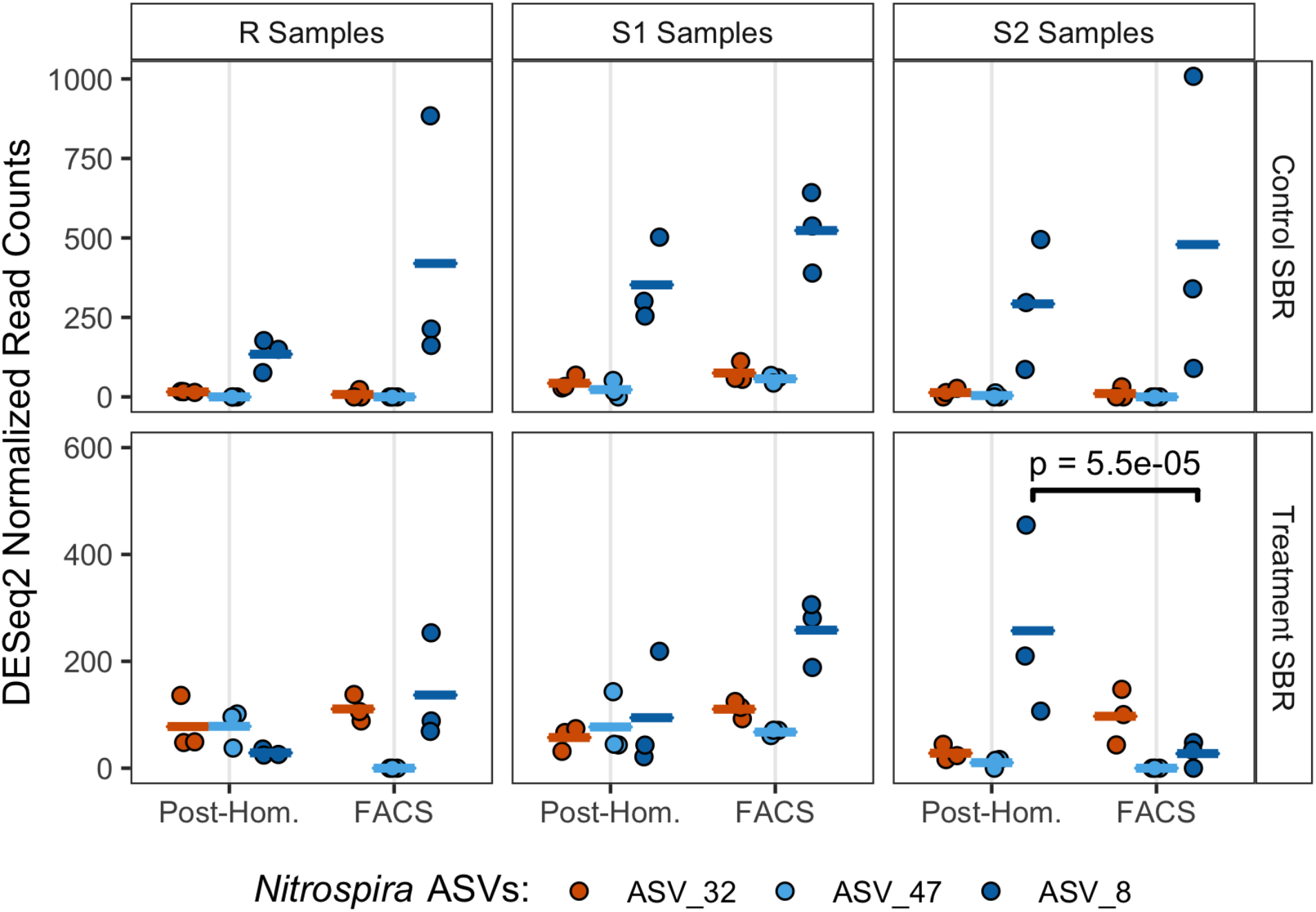
Normalized read counts of three dominant *Nitrospira* ASVs in triplicate BONCAT+ (i.e. FACS) and corresponding Post-Homogenized (i.e. Post-Hom.) and BONCAT+ (i.e. FACS) libraries generated from samples prepared from nitrifying microcosms seeded with mixed-liquor preceding return sludge treatment (R), return sludge after 15 min of side-stream treatment (S1), and return sludge after 24 h of side-stream treatment (S2) from the treatment SBR (bottom) and control SBR (top). Points indicate triplicate normalized read counts per ASV, and the horizontal bars of the same colors represent the sample mean per ASV. The reads were normalized with DESeq2 based on total read-counts per sample. A black bracket between an ASV within two samples represents a significant difference in the mean read abundance, determined with DESeq2 at an adjusted significance level of *p* < 0.01.

## Discussion

Implementing process control strategies that achieve consistent modulation of key functional groups remains a critical challenge towards the development of sustainable wastewater treatment biotechnologies. This is particularly true for repressing functionally diverse NOB to promote nitritation for energy-efficient biological nitrogen removal in AS processes. This study demonstrates how underlying shifts in the abundance of physiologically diverse *Nitrospira* populations can confer resilience to the nitrite-oxidizing community during an imposed press disturbance in AS treatment, in this case induced by routine exposure of return sludge to FA. Our results also demonstrate the utility of substrate analog probing approaches like BONCAT to illuminate the *in situ* ecophysiology of shared niches within the activated sludge microbiome, and the associated impacts on process and ecosystem functional stability.

We observed that routinely exposing ~20% of the return sludge to FA (200 mg NH_3_-N/L) in a sidestream reactor with a 24 hr retention time initially reduced nitrite-oxidizing activity in the mainstream reactor, achieving a maximum observed NAR of 42%. However, we observed acclimation of the nitrite-oxidation function after approximately 40 days of the press disturbance, indicated by decreasing NARs within the treatment SBR. This finding is in contrast to those reported by Wang et al. (21) who observed stable repression of NOB activity for over 100 days using a press disturbance of routine FA-exposure at similar conditions (210 mg N/L of FA for 24 hr, 22% return sludge exposed). It is important to note that both the pH of the side-stream sludge treatment incubation, and the daily input of total ammonia nitrogen into the mainstream SBR with the return sludge, were controlled across both SBRs. Furthermore, while salinity has been shown to be a driver of Nitrospira population structure (55), the salinity in the side-stream reactor during FA-exposure (~2.5 g/L as Na^+^ + Cl^-^) was 10-times less than the 50% inhibition level observed for a *Nitrospira* dominated AS community of 30 g/L NaCl (56). Therefore, the inhibition and acclimation of NOB activity to the applied press disturbance can likely be attributed to varying physiological responses of NOB community members to FA-exposure.

While acclimation of NOB communities in mainstream AS to routine FA-exposure has been previously reported (41, 42, 57), very few studies have directly investigated the role of physiological diversity between NOB members in supporting community-level acclimation. To our knowledge, all previous reports of NOB community acclimation to return sludge FA-exposure have involved shifts in major NOB genera during this press disturbance. Specifically, Li et al. (42) found that *Candidatus* Nitrotoga replaced *Nitrospira* as the dominant NOB in response to return-sludge FA-exposure, while Duan et al. (41) reported a shift from *Nitrospira* to *Nitrobacter* in response to FA-exposure. These reported shifts could be supported by potentially distinct physiological characteristics of *Candidatus* Nitrotoga and *Nitrobacter* compared to *Nitrospira,* such as higher tolerances to FA-inhibition (42, 58) and/or preferences for higher nitrite concentrations (30, 59). In contrast to these genus-level community shifts however, we observed that NOB community acclimation to FA-exposure could occur via shifts between *Nitrospira* sequence variants, specifically from a dominant variant belonging to *Nitrospira* lineage I (ASV_8) to two variants that belonged to *Nitrospira* lineage II (ASV_32 and ASV_47). These findings therefore reveal that physiological diversity at the lineage level, and possibly at the sub-lineage or strain level, within *Nitrospira*-dominated communities can facilitate niche-partitioning and acclimation to FA-exposure as a press-disturbance when applied as an engineering control strategy to promote low-energy nitrogen removal.

*Nitrospira* species are well-known to be metabolically versatile, with a wide range of functional potentials including nitrite oxidation, hydrogen oxidation, urea conversion, formate oxidation, nitrate reduction, and complete ammonium oxidation (24, 37, 38, 52, 60). This metabolic diversity likely creates opportunities for functionally degenerate *Nitrospira* populations to co-exist within a bioreactor by differentiating in niche space through auxiliary metabolic specializations, while also sharing a common niche space of nitrite oxidation (34, 61). Here, we employed BONCAT-FACS for the first time in an AS wastewater treatment system, to the best of our knowledge, which highlighted differing *in situ* physiologies of three *Nitrospira* variants and resolved their responses to FA-exposure on the level of cellular translational activity. BONCAT-FACS revealed that transitional activity in *Nitrospira* ASV_8 was significantly reduced following exposure to FA for 24 h, aligning with the results of the time-series reactor sampling in which *Nitrospira* ASV_8 was washed out of the treatment SBR but remained dominant in the control SBR. In contrast, *Nitrospira* ASV_32 and ASV_47 increased in abundance within the treatment SBR coinciding with the decrease in NAR after day 40, and BONCAT-FACS revealed the translational activity of those variants remained unchanged following FA-exposure. Based on the close alignment of the BONCAT-FACS observations with the trends in the time-series reactor sampling, it can therefore be hypothesized that the wash-out of *Nitrospira* ASV_8 occurred due to its decreased activity in response to the press-disturbance of routine sidestream FA-exposure, which induced a growth lag once it was reintroduced into the mainstream SBR. In contrast, the potentially distinct auxiliary metabolic potentials of *Nitrospira* ASV_32 and ASV_47 may have conferred their physiological resistance to the side-stream FA-exposure, thereby providing these variants with a competitive growth advantage within the mainstream SBR. Physiological tolerance to FA-exposure was identified as a mechanism of niche-partitioning between *Nitrospira* populations of lineages I and II by Ushiki et al. (40), who showed that *Nitrospira* sp. ND1 of lineage I was more sensitive to FA compared to *Nitrospira japonica* of lineage II at 100 mg NH_4_^+^-N/L and pH 8.0. Here, we observed that *Nitrospira* ASV_32 and ASV_47 of lineage I were physiologically more tolerant to FA compared to *Nitrospira* ASV_8 of lineage II, suggesting that either lineage-specific tolerances are distinct at the higher FA concentration applied here (200 mg NH_3_-N/L) or that physiological tolerance to FA is a trait that varies at the strain/species level within *Nitrospira.* Regardless, once *Nitrospira* ASV_32 and ASV_47 grew to high enough abundances in the treatment SBR, they likely contributed to the net oxidation of nitrite and the decrease in the NAR that was observed after day 40 of the Treatment Phase. Therefore, the FA-exposure press disturbance strategy employed in this study acted as a selective pressure that impacted the stability of the NOB community composition, and functional degeneracy within the *Nitrospira* sequence variants likely provided the nitrite-oxidizing community with resilience by shifting activity to physiologically resistant community members.

The above results highlight the need to combine press disturbances that target the distinct physiological traits of multiple functionally degenerate NOB populations, defined at both the inter and intra genus level, to effectively reduce the aggregate activity of the NOB community for energy-efficient nitrogen removal. For example, if future studies support our finding that lineage I *Nitrospira* are more tolerant to FA than those of lineage II, then combining press-disturbances that further target the physiological traits of lineage I *Nitrospira* along with FA-exposure could prove efficacious. Such strategies could entail the maintenance of high dissolved oxygen concentrations, based on adaptation of lineage I *Nitrospira* to low dissolved oxygen environments (34, 36), or the maintenance of low nitrite concentrations based on their preference for higher nitrite concentrations (34, 40, 51). Measurements of maximal growth rates for these sequence variants could also inform wash-out strategies based on limiting the solids retention time (SRT) (13). Acknowledging the potentially tight tolerances and dynamic nature of nitrite, oxygen, and SRT controls required to target the biokinetics of lineage I *Nitrospira* in full-scale AS systems, combinatorial press-disturbance strategies would likely benefit from control strategies that incorporate frequent community monitoring and biokinetic data of these physiologically diverse NOB populations. These findings also underscore the need for broader applications of *in situ* physiology approaches for elucidating the impacts of NOB out-selection strategies on functionally active NOB members within activated sludge systems.

Accurately measuring the active biomass fraction in microbial bioprocesses is critical, as many key biokinetic models and process mass balances are based on active biomass concentrations (62). However, conventional indirect approaches for quantifying active biomass based on net substrate utilization and growth yields are unable to resolve the compositional and functional dynamics of active microbial populations. As enzymes are the key catalysts that drive the majority of substrate transformations in microbial bioprocesses, we posit that measures of active biomass should be based on the translationally active microbial cell fraction. The translationally active cell fractions in both SBR microcosms seeded with mixed liquor measured by BONCAT-FACS (R; 24.1% ± 2.6% in treatment, 30.1% ± 3.8% in control) were less than the active biomass fraction predicted through steady-state modeling (63) (67-77% VSS-basis; Supplemental Information), which is likely because the microcosms were only supplemented with ammonium as an electron donor. Nonetheless, we observed many translationally active heterotrophs in the BONCAT microcosms, which could have remained active through metabolism of internal carbon reserves, endogenous respiration, or constitutive protein expression. The consistent increase in translational activity across all differentially abundant ASVs detected in the BONCAT+ fraction of the control SBR microcosm seeded with sludge following the 24 h sidestream incubation could thus have been attributed to fermentative metabolism on cell decay products. In contrast, the mixed translational responses of differentially abundant ASVs in the BONCAT+ fraction of the treatment SBR microcosm following sidestream incubation with FA suggests that more complex dynamics between cellular inhibition, decay, and fermentative metabolism were induced by FA-exposure. This analysis highlights the value of BONCAT as an *in situ* physiology approach to directly measure concentrations of translationally active population members in a microbial bioprocess. This approach could therefore be extended to provide actual measurements of the active biomass concentration for use in calibrating and validating higher-resolution process models, which have been called for as tools to optimize energy-efficient nitrogen removal technologies (9) as well as to better resolve biogeochemical cycles in natural ecosystems (64, 65). BONCAT-FACS could also be extended to resolve associations in the activity of AS taxa with different substrate preferences (48), thus helping to validate ecological-scale models of the wastewater microbiome (66). Therefore, *in situ* physiology approaches like BONCAT show great promise to help inform new strategies to model, control, and engineer microbiome function in both environmental biotechnologies and natural ecosystems alike.

## Materials and Methods

### Reactor setup, operation and monitoring

Two identical lab-scale sequencing batch reactors (SBRs) with working volumes of 4.28 L were seeded with AS from a pilot-scale SBR at a WWTP in King County (Washington, USA) (Figure S1). The SBR cycles lasted 3 h, including 2 min aerobic feeding, 148 min aerobic reaction, 20 min settling, 5 min decanting, and 5 min idle periods. The SBR cycle timing was controlled with ChronTrol XT timers (ChronTrol Corporation, USA), and mixing was provided by overhead mixers. Reactor temperature was maintained at 20 ± 1°C using an environmental chamber. The target solid retention times (SRTs) were kept at 10 days for both SBRs throughout the study by wasting a determined amount of biomass based on daily values of total suspended solids (TSS) and volatile suspended solids (VSS) measured in the mixed liquor and effluent streams. The SBRs were fed with a synthetic wastewater containing ammonium chloride as the nitrogen source and sodium acetate and propionic acid as organic carbon sources, producing 24.5 ± 2 mg NH_4_^+^-N/L and 100 ± 37 mg/L of soluble chemical oxygen demand (sCOD), respectively, to reflect a sCOD/NH_4_^+^-N ratio of ~4.0 typical of North American wastewaters (63). Macro and micro element components of the synthetic wastewater are detailed in the Supplemental Information.

Two operational phases were sequentially conducted: the Start-up Phase and the Treatment Phase. In the Start-up Phase, the two SBRs were operated under the same aerobic conditions without FA-exposure of return sludge to achieve similar nitrification performance. The SBRs were fed with 1.07 L of synthetic wastewater in each SBR cycle using peristaltic pumps (LabF1/YZ1515, Shenchen Precision Pump, China), resulting in a hydraulic retention time (HRT) of 12 h. The pH was not controlled but measured within the range of 6.7-7.5. Aeration was provided by an air pump and dissolved oxygen was not controlled, but ranged from 3-8 mg/L in a typical SBR cycle for both reactors. The Start-up Phase lasted 274 days to establish steady-state conditions.

The Treatment Phase was commenced on day 275, hereafter referred to as ‘day zero’, and lasted 94 days. The operational conditions in the Treatment Phase were similar to those in the Start-up Phase, except for the following differences (Figure S1). In the Treatment Phase, 800 mL of mixed liquor was removed from each SBR at the end of the reaction period of a given cycle every 24 h, and was thickened to 50 mL by centrifugation. The thickened return sludge (50 mL) was incubated in an unstirred 200 mL beaker along with 100 mL of media with the same composition as the synthetic feed, but with no COD. For the experimental reactor, termed the treatment SBR, the return sludge treatment solution contained 1060 mg/L NH_4_^+^-N, with pH adjusted to 9.0 using sodium hydroxide to produce a final FA concentration of 200 mg NH_3_-N/L. These FA and total ammonia nitrogen concentrations are within the range observed in anaerobic digester centrate, particularly co-digestors and those with thermal-hydrolysis pre-treatment (67, 68). In the other reactor, termed the control SBR, thickened return sludge was incubated in the same medium at a pH of 9.0 but without ammonium addition. After 24 h of incubation, the 150 mL of treated return sludge was recycled back into the respective SBRs at the start of the next cycle. To maintain consistent nitrogen loadings in the two SBRs, 0.406 g of ammonium chloride was added to the control SBR simultaneously with the treated return sludge. Monitoring experiments lasting 24 h were performed approximately every 10 days to measure composite daily nitrite accumulation ratios (NARs), as described in the Supplemental Information.

During the Treatment Phase, effluent nutrient samples were collected just before the addition of treated return sludge. Nitrogen species (ammonium, nitrite, and nitrate), orthophosphate, and soluble COD (sCOD) in effluent samples were monitored three to four times per week. pH and DO were measured at least three times per week. Analytical methods for bioreactor monitoring are described in Table S2.

### Microcosms for bioorthogonal non-canonical amino acid tagging (BONCAT)

Three 30 mL samples were collected from each SBR throughout the return sludge treatment cycle commenced on day 94, including: (1) a mixed liquor sample immediately preceding bulk mixed liquor removal for return sludge treatment; (2) a return sludge sample after 15 min of side-stream treatment (S1); and (3) a return sludge sample after 24 h of side-stream treatment (S2). The treated return sludge samples were volume-corrected for the thickening process to attain the same biomass concentrations as the mixed liquor samples. Samples were washed in phosphate buffered saline (1x PBS; filter sterilized) through centrifugation (3000 rpm, 5 min) and resuspension to remove residual growth substrates. All samples were incubated in a media consisting of 25% synthetic wastewater in 1x PBS (vol/vol) without COD or yeast extract, so that NH_4_^+^-N was the only exogenous electron donor. For BONCAT-labeling microcosms, 15 mL portions of each sample were resuspended in 15 mL of incubation media amended with 1 mM homopropargylglycine (HPG; Click Chemistry Tools, USA) (HPG-positive microcosm), transferred into sterile 125 mL Erlenmeyer flasks, and incubated on an orbital shaker for 3 h at 20 °C and 200 rpm. Control microcosms (HPG-negative) for each sample were conducted similarly except without HPG amendment. Following incubations, microcosm samples were washed three times in 1x PBS to remove unincorporated HPG, resuspended in 10% (vol/vol) glycerol in PBS, aliquoted into 1 mL and stored at −80°C until further processing. Details on preliminary BONCAT validation microcosms are provided in the Supplemental Information.

### BONCAT Sample Preparation and Click-Chemistry

For each microcosm type (R, S1, or S2) for both SBRs, sample preparation and click-chemistry were conducted using triplicate HPG-positive and duplicate HPG-negative microcosm samples. Samples were thawed on ice at 4°C, enzymatically homogenized, subjected to filter-immobilized click-chemistry labeling with the FAM picolyl azide dye (Click Chemistry Tools, USA) closely following the procedure of Couradeau et al. (45), detached from the filter, pre-strained through a 30 um mesh filter, and counterstained with SYTO59 (10,000x final dilution; ThermoFisher Scientific, USA) to generate the pre-sort samples for FACS. Details of the sample homogenization and click-chemistry labeling procedures are provided in the Supplemental Information.

### Fluorescence-activated cell sorting (FACS)

Fluorescence-activated cell sorting was conducted on a BD FACSJazz™ cell sorter (BD Biosciences, USA) calibrated to detect the FAM picolyl azide dye (excitation = 490 nm/emission = 510 nm) and the SYTO59 counterstain dye (excitation = 622 nm/emission = 645 nm). An overview of the FACS gating procedures is provided in the Supplemental Information. Briefly, initial gating was established with side-scatter, forward-scatter and trigger pulse width to exclude large particles and cell aggregates. Sorting gates were set targeting BONCAT-positive (BONCAT+) cell fractions based on background SYTO59 and FAM fluorescence, allowing less than 0.5% of false positives (Figures S13-S14). A total of 100k cells were analyzed from each sample, where cells within the sorting gate (SYTO+, BONCAT+) were sorted into 1.5 mL tubes containing 400 μL of prepGEM Wash Buffer (ZyGEM™, USA), and stored at −80 °C until further analysis.

### DNA Extraction and 16S rRNA gene amplicon sequencing

For time-series analysis of community composition, triplicate 980 μL aliquots of mixed liquor were routinely collected directly from the SBRs at the end of a cycle and flash frozen at −80°C. Mixed liquor samples were thawed on ice, and DNA was extracted using the FastDNA Spin Kit for Soil (MP Biomedicals, USA) with minor modifications (69). For each HPG-positive microcosm type (R, S1, or S2) for both SBRs, 50 μL aliquots were collected from the triplicate pre-homogenized and post-homogenized samples during preparation for FACS, added to 400 μL of prepGEM Wash Buffer, and stored at −80°C. Pre-homogenized samples from HPG-negative R microcosms and bulk mixed liquor samples were prepared similarly. All pre-homogenized, post-homogenized, and BONCAT-FACS samples were extracted using the prepGEM Bacteria kit (ZyGEM™) using a low-biomass input procedure overviewed in the Supplementary Information. DNA concentrations were measured with the Qubit^®^ dsDNA BR and HS Assay Kits and Qubit^®^ fluorometer (ThermoFisher Scientific, USA).

16S rRNA gene fragments from all DNA extracts were amplified using barcoded primers, 515F and 926R (70), targeting the V4-V5 hypervariable region of the 16S rRNA gene. All samples extracted with the prepGEM kit underwent an initial round of 15-cycles of PCR with non-barcoded 515F and 926R primers due to the low-biomass input. One triplicate set of FastDNA extracts from day 94 were also pre-amplified to determine potential biases of that step. Amplified barcoded PCR products were pooled at equimolar concentrations and sequenced on an Illumina MiSeq in paired-end 300 bp mode at the UBC Biofactorial facility.

Amplicon reads were processed and denoised into amplicon sequencing variants (ASVs) with DADA2 (49) (v.1.12.1) in the R environment. The script used to generate the ASV datasets is provided in Supplemental Information. Denoised sequences were taxonomically classified using the RDB Classifier (71) against the MiDAS 3.0 database (69).

### Statistical analysis

Comparisons of reactor nutrient data were performed with t-tests for N-of-1 trials with serial correlation (72). Amplicon sequencing data was visualized using the tidyverse package (73) (v.1.3.0) in the R environment. Principal coordinates analysis (PCoA) of cumulative-sum scaled (CSS) ASV read counts was performed using the metagenomeSeq (74) (v.1.26.3) and vegan (75) (v.2.5.6) packages in R. PERMANOVA (adonis) was conducted in vegan with 1000 permutations to determine significant differences in reactor communities over time. Differential abundance analysis of ASVs between SBRs and in BONCAT microcosm datasets was performed using DESeq2 (76) (v.1.24.0) using parametric fitting, the Wald significance test, and Benjamin-Hochberg correction for *p* values. A log_2_-fold change of an ASV represents the multiplicative effect size for changes in normalized read counts across treatments on a logarithmic scale to base 2. Sequence similarity values between ASVs were calculated using the NCBI Basic Local Alignment Search Tool (77). FACS data was processed using the flowcore package (v.1.52.1) (78) in R.

## Data Availability

The raw read files of 16S rRNA gene amplicons are available via the NCBI Short Read Archive under BioProject PRJNA693634.

## Acknowledgements

This work was supported by the Natural Sciences and Engineering Research Council (NSERC) of Canada (Discovery Grant). MM was supported by an NSERC CGS-M fellowship.

## Author Information

MM and YL contributed equally to this work.

