## Supplemental Information for "Activity-based cell sorting reveals resistance of functionally degenerate *Nitrospira* during a press disturbance in nitrifying activated sludge"

[+] Contributed equally to this work.

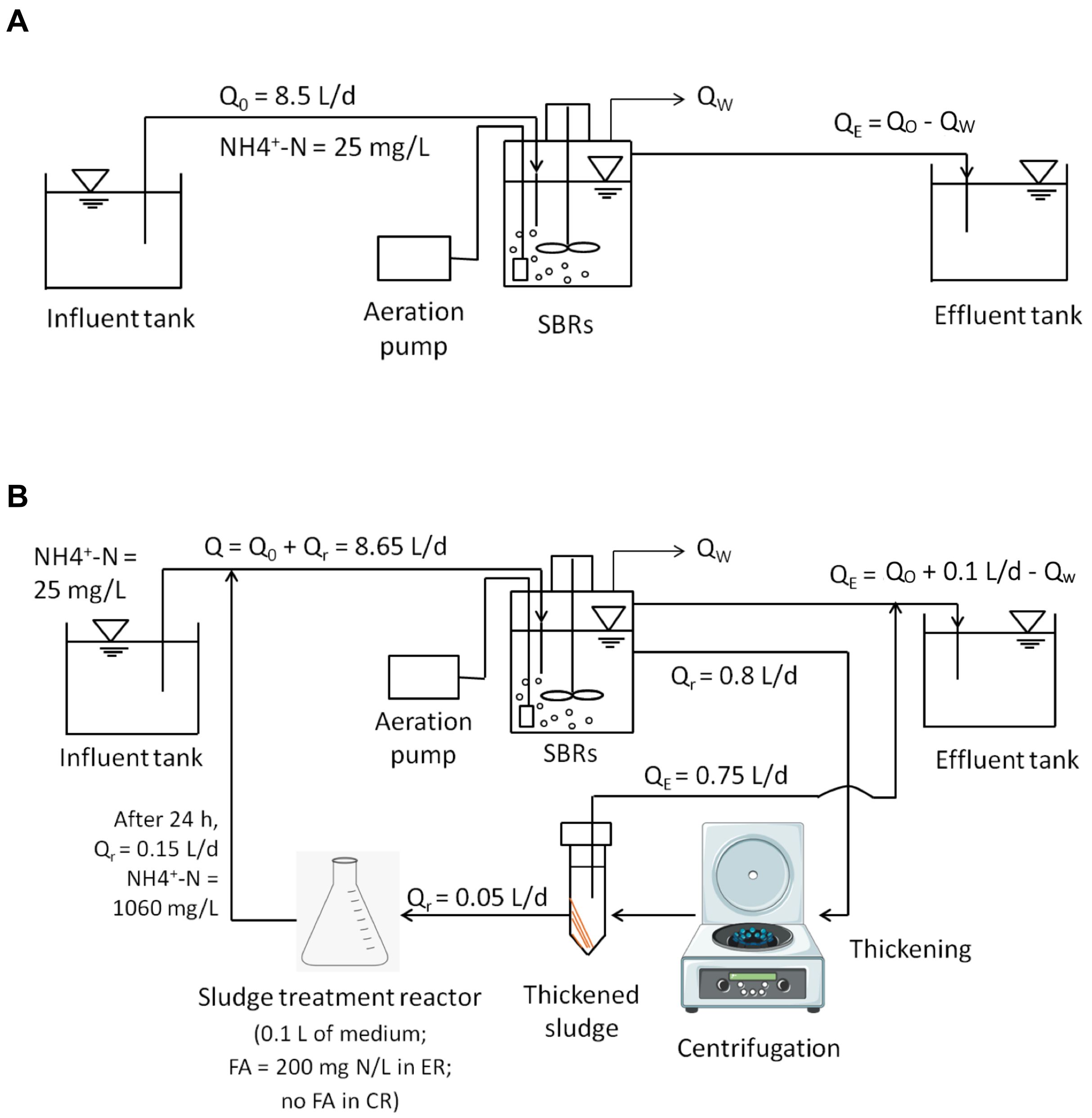

**Figure S1:** Schematic diagrams showing the operation of the two SBRs during (A) Start-up Phase and (B) Treatment Phase. ER = Experimental Reactor (also termed Treatment SBR); CR = Control SBR; FA = free ammonia (mg $NH_3$-N/L); $Q_0$ = feed flow rate of synthetic wastewater; $Q_E$ = effluent flow rate; $Q_W$ = waste activated sludge flow rate; $Q_r$ = side-stream return sludge treatment flow rate.

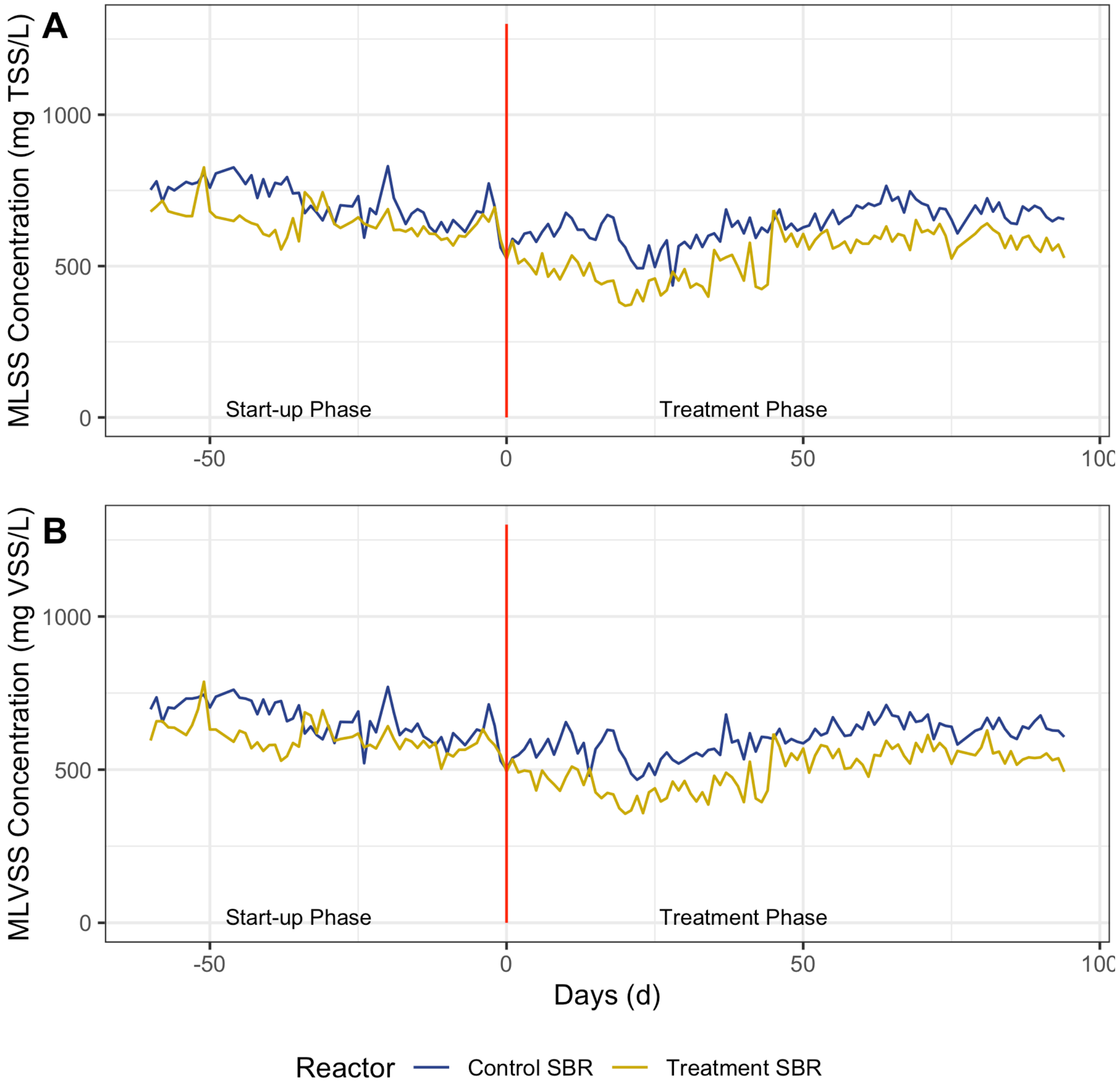

**Figure S2:** (A) Mixed liquor total suspended solids (MLSS) and (B) Mixed liquor volatile suspended solids (MLVSS) for both SBRs in the Start-up and Treatment operational phases.

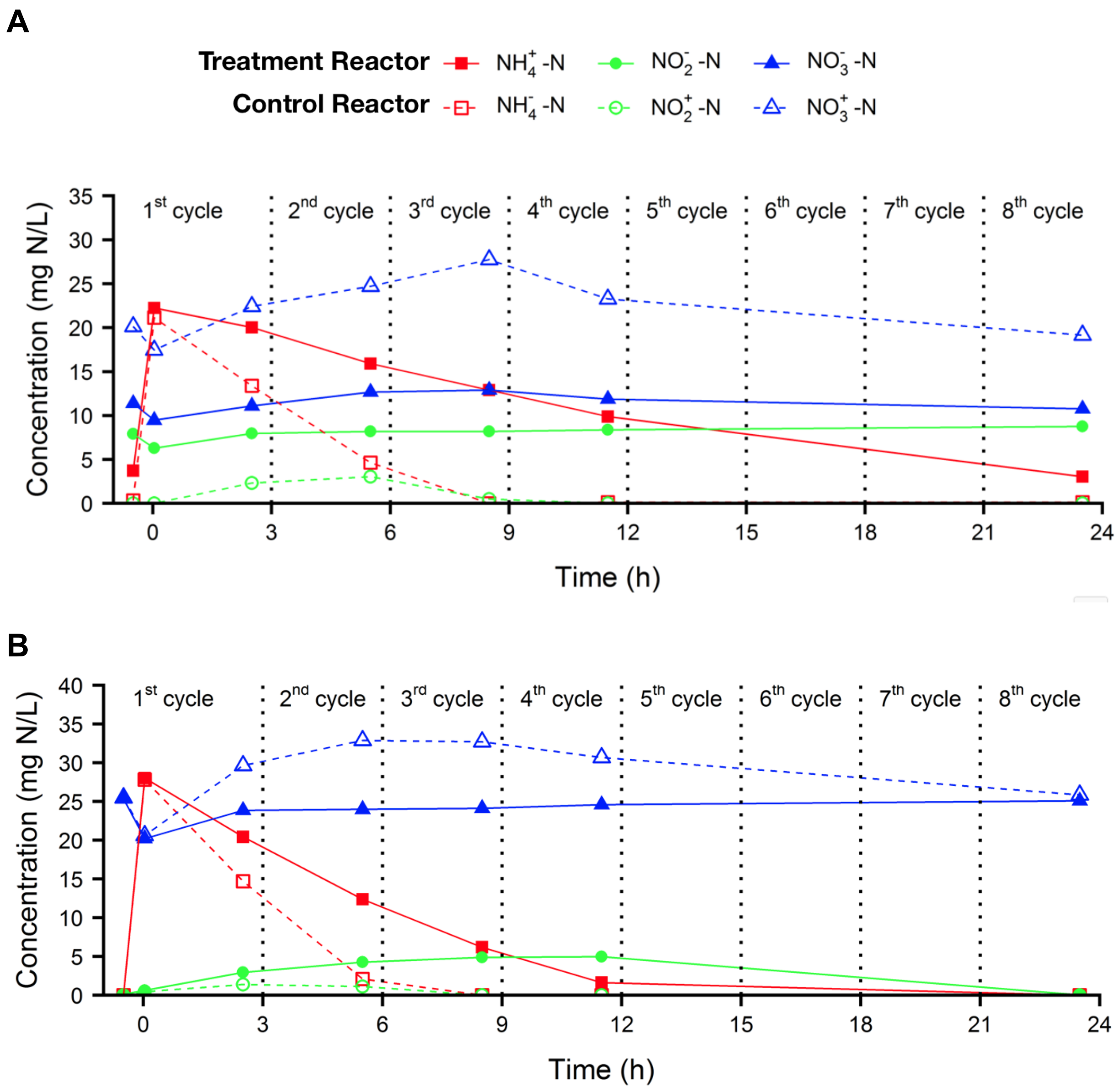

**Figure S3:** Ammonium ($NH_4^+$-N), nitrite ($NO_2^-$-N), and nitrate ($NO_3^-$-N) concentrations in the treatment SBR and control SBR throughout a 24 h period, measured on days (A) 37; and (B) 75 of the Treatment Phase. The timing of the SBR cycles are marked with vertical dashed lines.

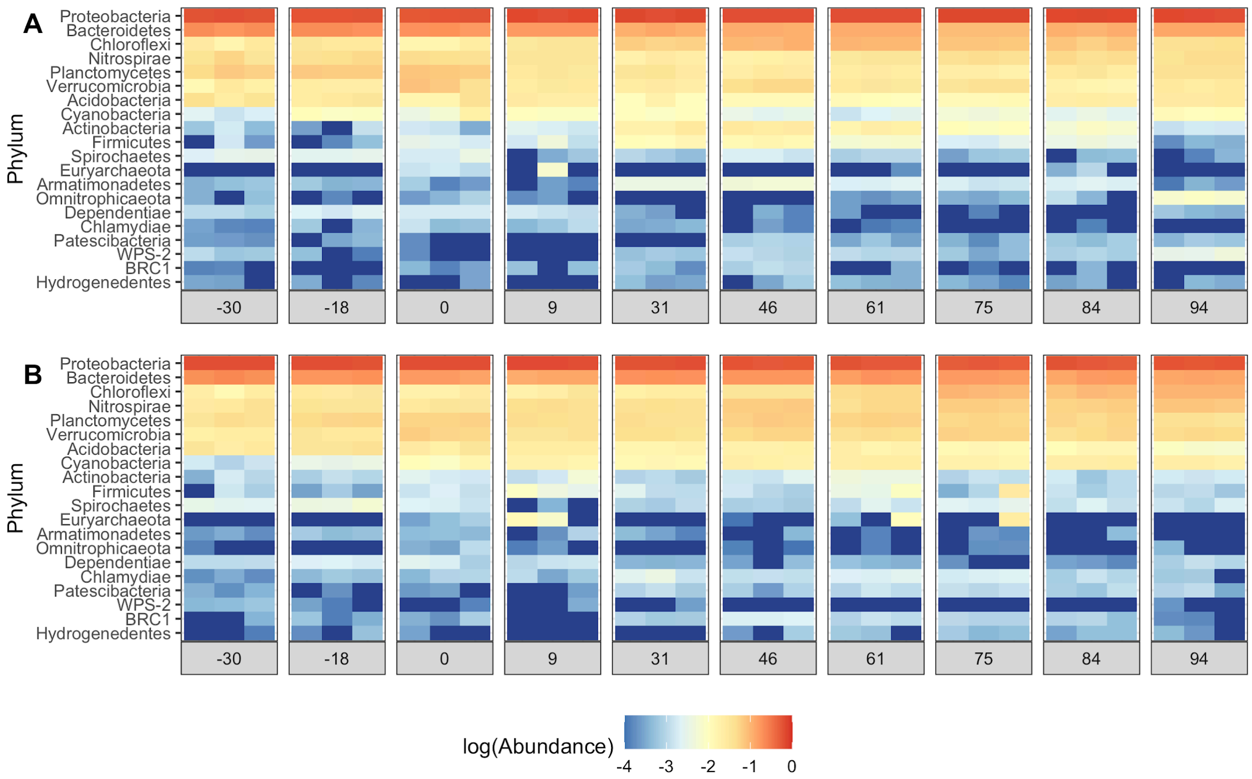

**Figure S4:** Heatmap of log-scaled relative abundance (total sum scaled) of top 20 most abundant phyla in the (A) treatment SBR and (B) control SBR over the two operational phases. Sequencing results are shown for triplicate DNA extractions on each sampling date. Phyla are ordered from highest to lowest cumulative abundance across all samples.

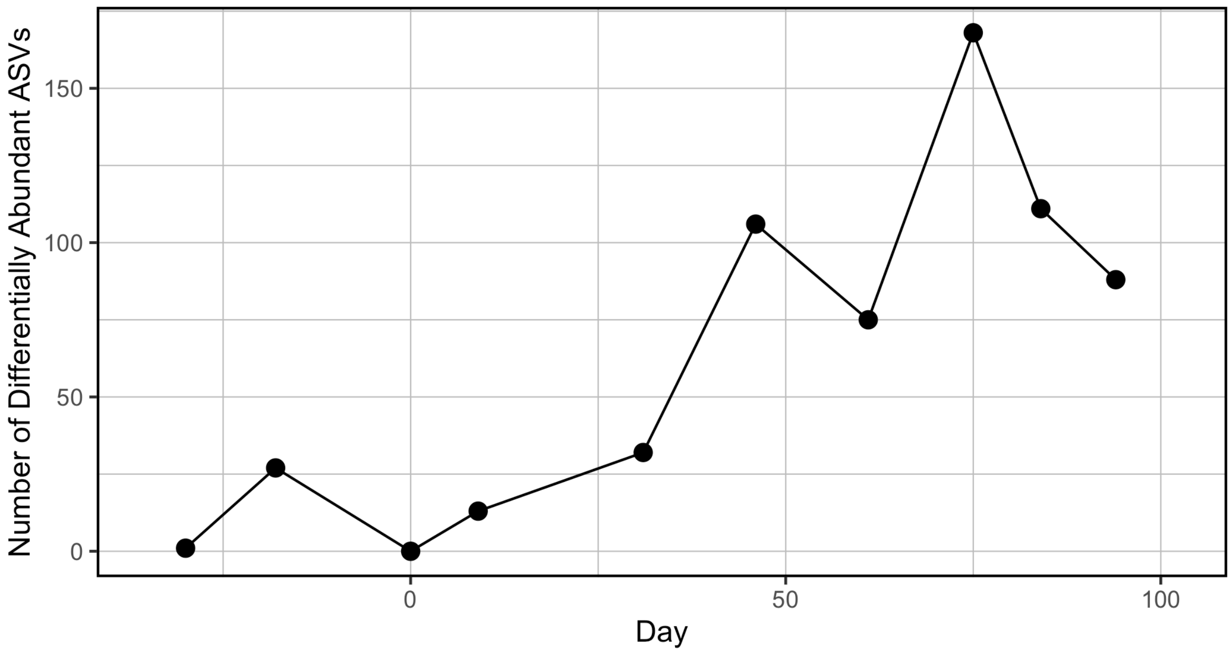

**Figure S5:** Number of differentially abundant ASVs between the treatment and control SBRs over the two operational phases, as determined by comparing ASV abundance in triplicate DNA extracts at each timepoint using DESeq2 v.1.24.0 with a significance level of $p < 0.01$.

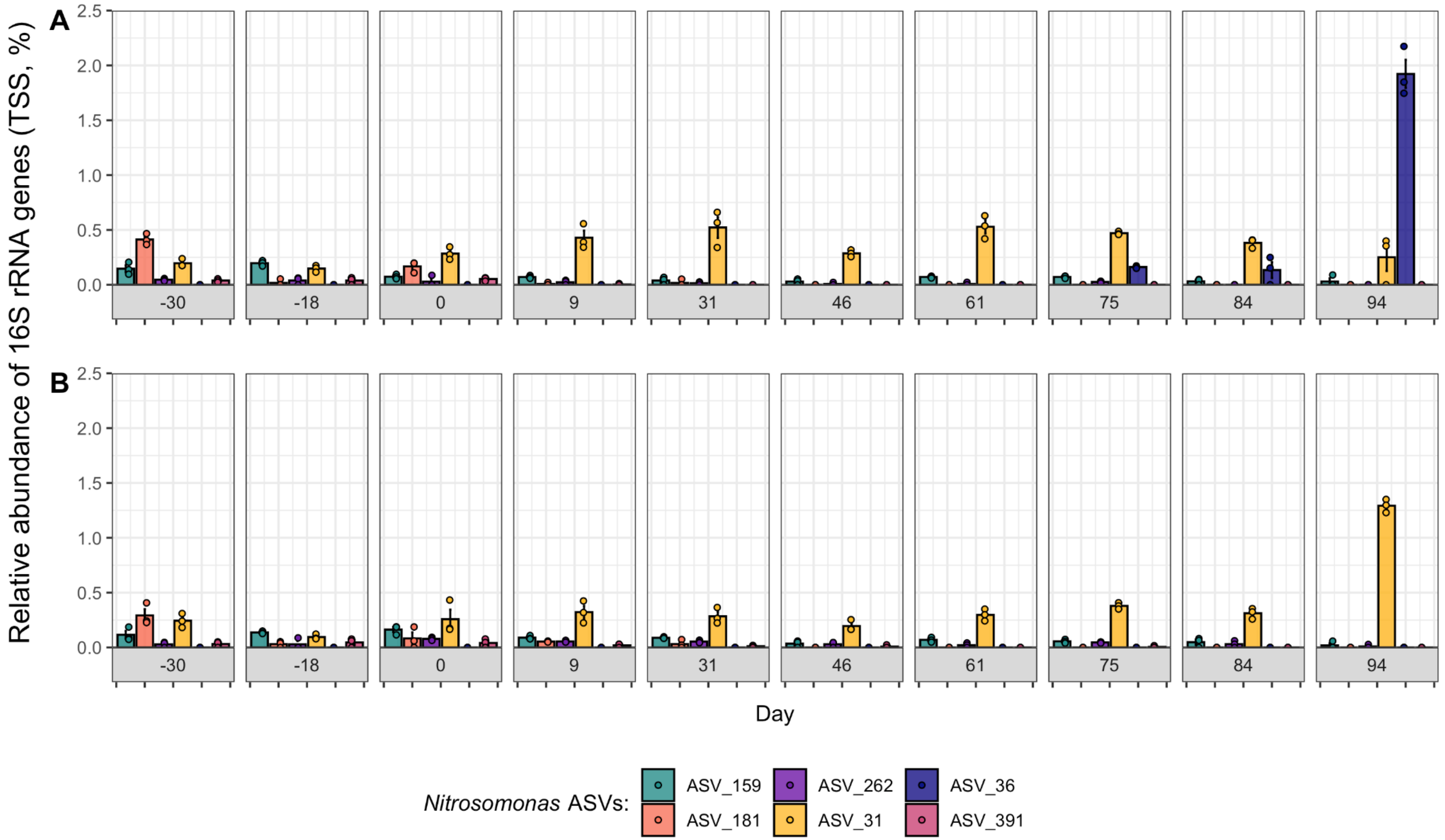

**Figure S6:** Relative abundance (total sum scaled, TSS) of 6 dominant *Nitrosomonas* ASVs over time in the (A) treatment SBR and (B) control SBR. Only *Nitrosomonas* ASVs with mean relative abundances greater than 0.05% detected in at least one sample are shown. Results are shown for triplicate DNA extractions on each day, with the coloured points showing the relative abundance of each ASV in each DNA extraction, the coloured bar showing the mean relative abundance, and the error bars showing the standard error of the mean.

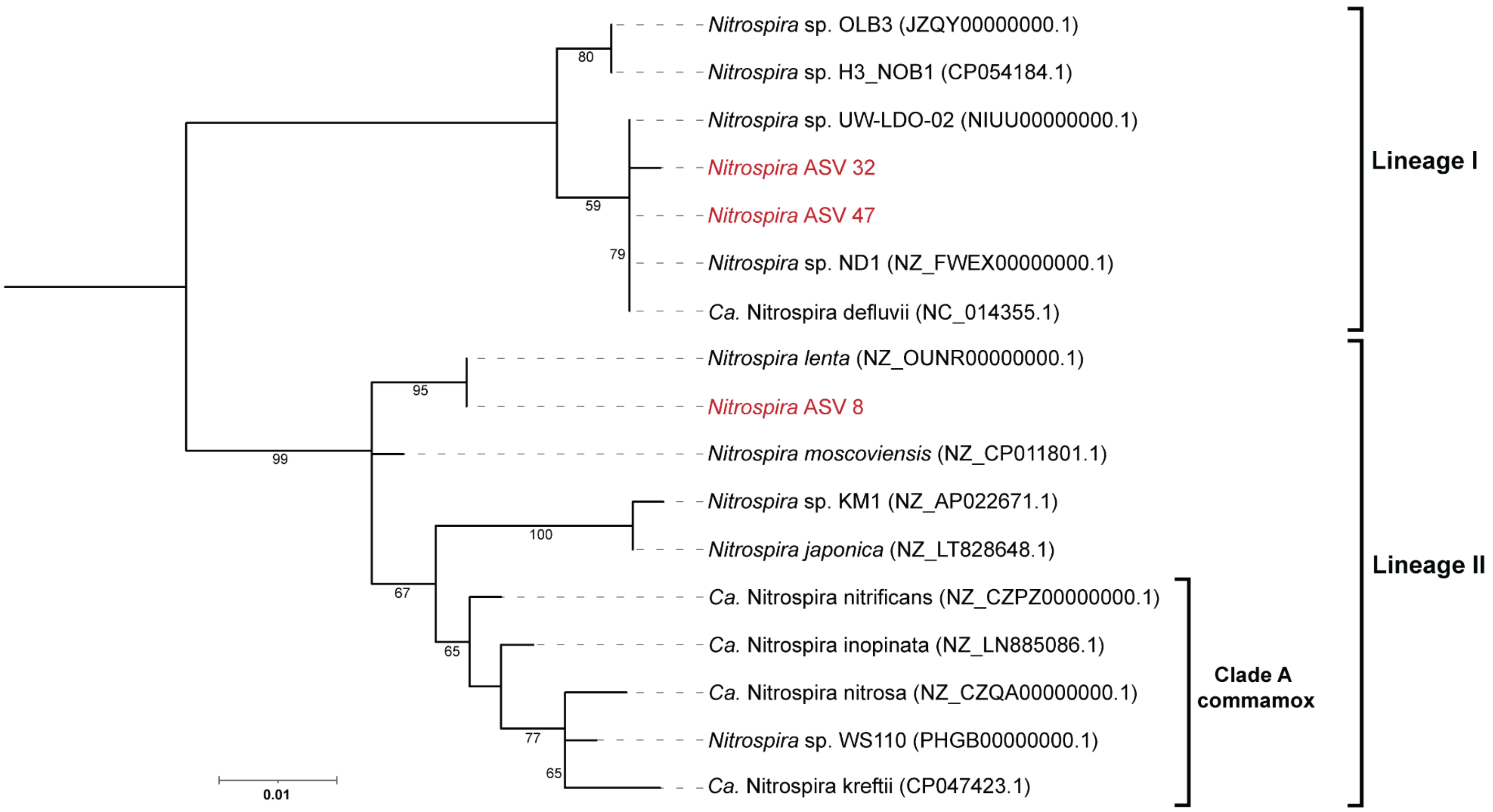

**Figure S7:** Phylogenetic analysis of partial 16S rRNA gene sequences of the *Nitrospira* ASVs detected in this study (ASV_8, ASV_32 and ASV_47; shown in red) in comparison to reference sequences of related *Nitrospira* species obtained from NCBI. To construct the phylogenetic tree, barrnap v0.9 (1) was first used to extract the 16s rRNA gene sequences from the full or partial NCBI reference genomes. The extracted reference 16S rRNA genes were then aligned to the partial 16S rRNA sequences of the three *Nitrospira* ASVs obtained in this study using MUSCLE (2), and the multiple sequence alignment was trimmed in MEGAX (3). The phylogenetic tree was constructed using iq-tree (4) with default parameters and 1000 non-parametric bootstrap replicates, and then visualized using iTOL (5). The maximum likelihood tree shows that *Nitrospira* ASV_8 was most closely related to *Nitrospira* lenta and other *Nitrospira* lineage II species, while ASV_32 and ASV_47 were more closely related to *Nitrospira* lineage I species including *Nitrospira* sp. ND1 and *Candidatus* Nitrospira defluvii. All *Nitrospira* ASVs detected in this study clustered away from the Clade A commamox *Nitrospira* species which have been identified in wastewater treatment plants. Branch node numbers represent bootstrap support values greater than 50%. Genbank accession numbers for the reference *Nitrospira* species are provided in parenthesis.

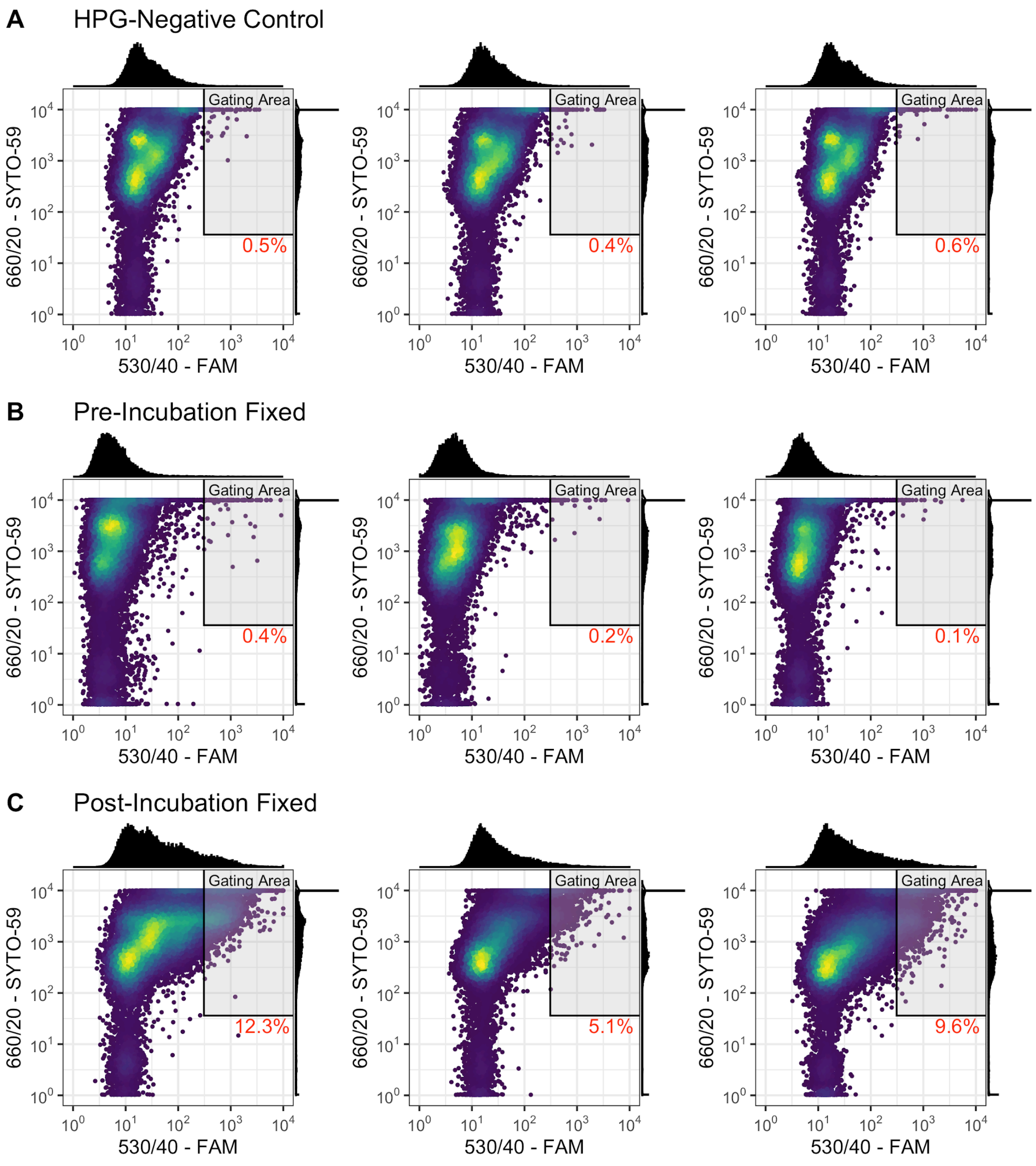

**Figure S8:** BONCAT-FACS results generated from preliminary validation microcosms in which cells were (A) incubated without HPG, (B) fixed with 3% paraformaldehyde before incubation with HPG, and (C) fixed with 3% paraformaldehyde after incubation with HPG (see Text S1). The gating area shows STYO+ and BONCAT+ cell fluorescence determination based on 0.5% false positives. The fractional abundance of BONCAT+ cells in each sample was calculated as the fraction of SYTO+ cells residing in the sorting gate, as indicated in red text in each box. The absence of BONCAT+ cells in the Pre-Incubated Fixed control samples (B), but detection of BONCAT+ cells in the Post-Incubation Fixed samples (C), indicates that cells needed to actively uptake and incorporate HPG to become fluorescently labelled by the FAM picolyl azide BONCAT dye. The lower detection rate of BONCAT+ cells within the Post-Incubation Fixed samples may be attributed to a potentially lower level of translational activity within the control SBR mixed liquor during the time of conducting the validation microcosms, or to potential impacts of cellular fixation on sample homogenization and/or click-chemistry labeling efficiencies.

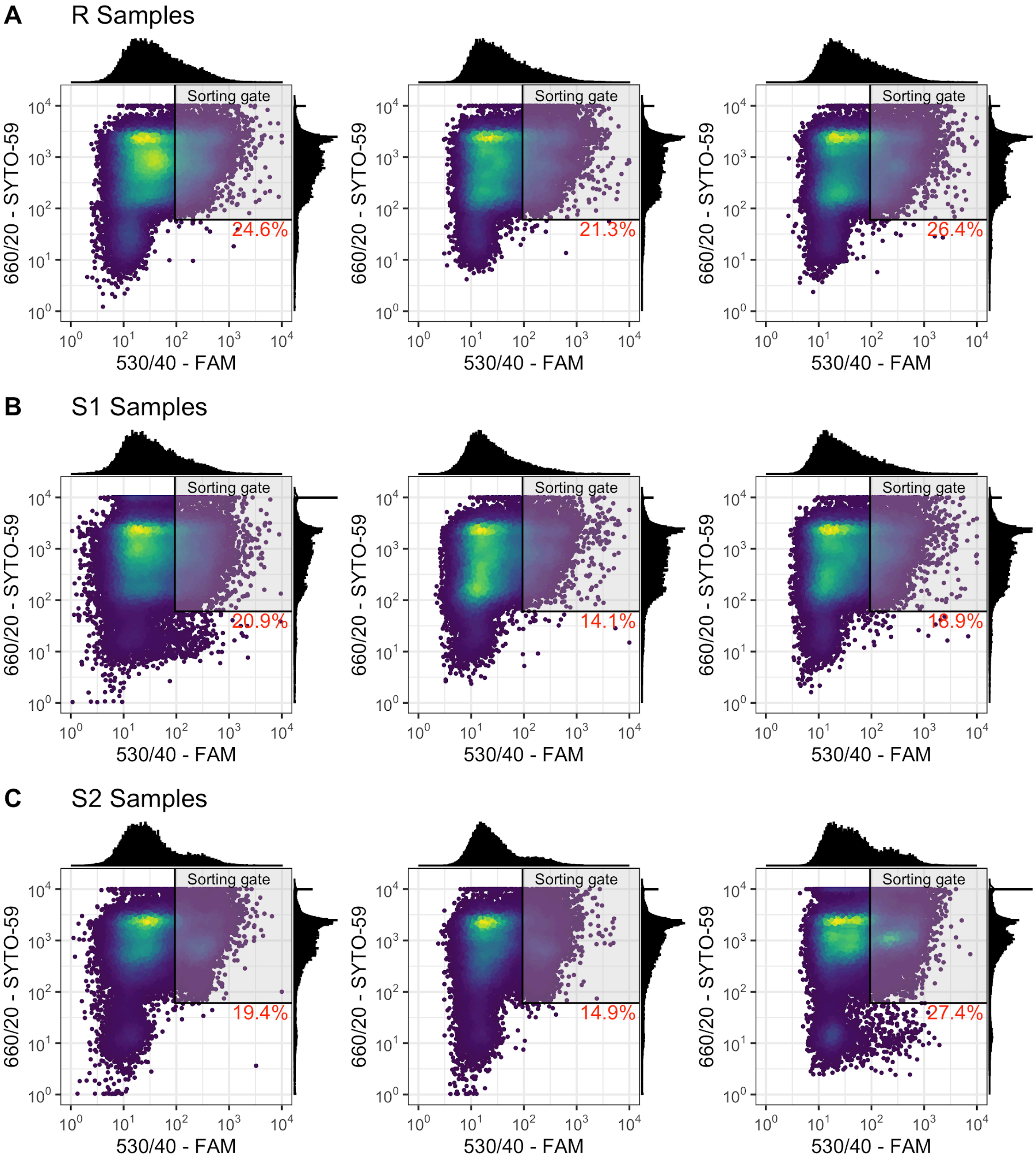

**Figure S9:** BONCAT-FACS results generated from treatment SBR microcosm samples. BONCAT-FACS was conducted on Post-Homogenized samples prepared from nitrifying microcosms seeded with the treatment SBR mixed liquor preceding return sludge treatment (R), return sludge after 15 min of side-stream treatment (S1), and return sludge after 24 h of side-stream treatment (S2). BONCAT+ cells residing in the predetermined (Fig. S2) sorting gate were sorted and collected from each sample for 16S rRNA gene amplicon sequencing. The fractional abundance of BONCAT+ cells in each sample was calculated as the fraction of SYTO+ cells residing in the sorting gate, as indicated in red text in each box.

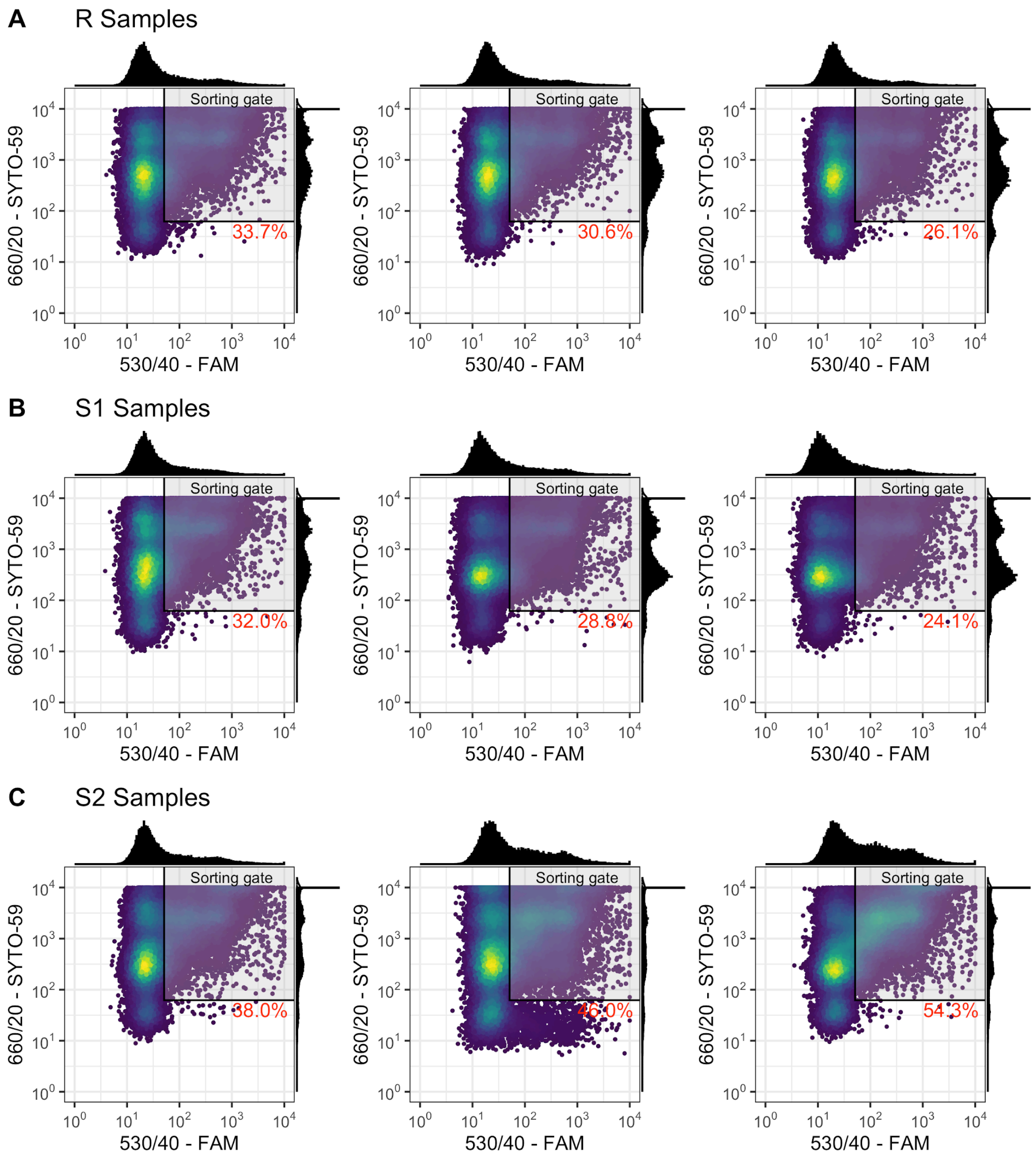

**Figure S10:** BONCAT-FACS results generated from control SBR microcosm samples. BONCAT-FACS was conducted on Post-Homogenized samples prepared from nitrifying microcosms seeded with the control SBR mixed liquor preceding return sludge treatment (R), return sludge after 15 min of side-stream treatment (S1), and return sludge after 24 h of side-stream treatment (S2). BONCAT+ cells residing in the predetermined (Fig. S3) sorting gate were sorted and collected from each sample for 16S rRNA gene amplicon sequencing. The fractional abundance of BONCAT+ cells in each sample was calculated as the fraction of SYTO+ cells residing in the sorting gate, as indicated in red text in each box.

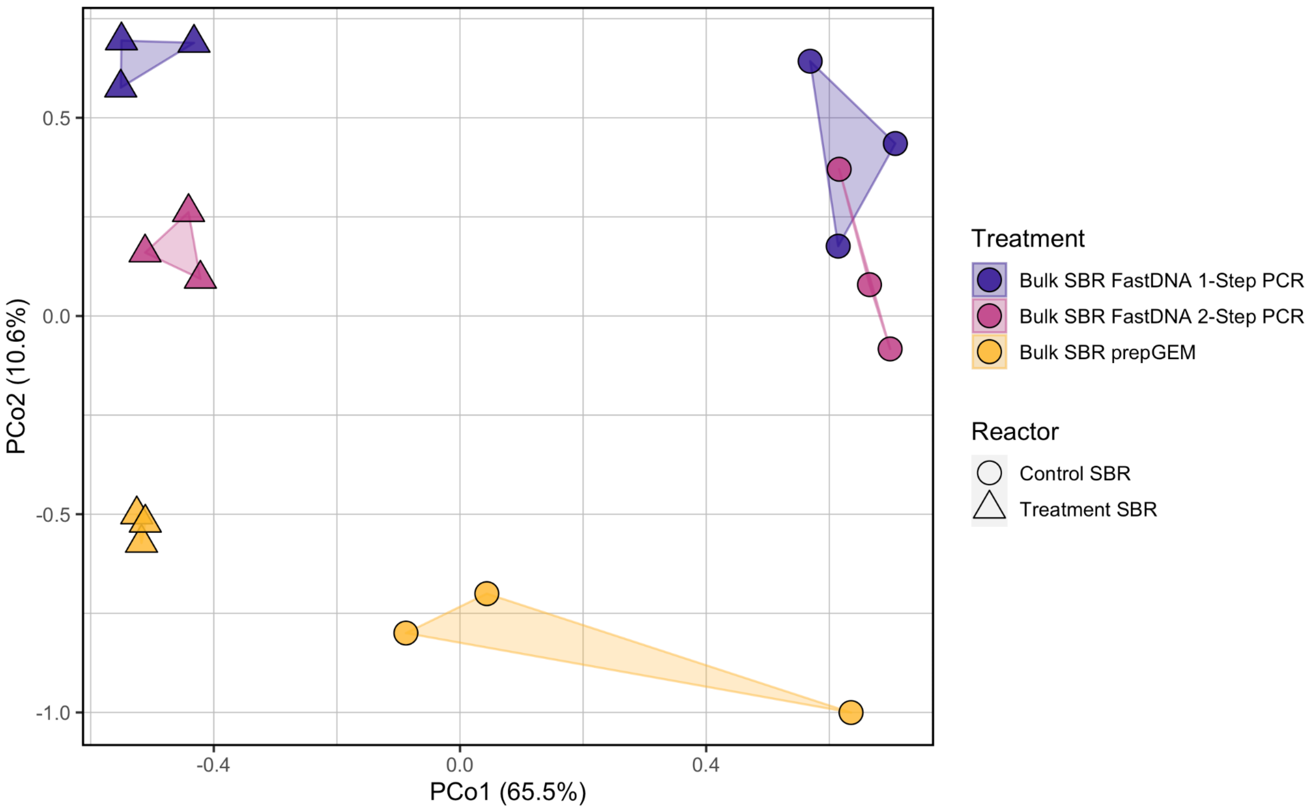

**Figure S11:** Principal coordinates analysis of cumulative sum scaled (CSS) read counts of 16S rRNA ASVs (V4-V5 region) measured in samples from the bulk mixed liquors of the control and treatment SBRs that were prepared using different DNA extraction kits (FastDNA Soil Kit or prepGEM Bacteria Kit; both amplified with 2-step PCR) and PCR amplification procedures (1-step or 2-step PCR; both extracted with FastDNA Soil Kit). The marker fill represents the origin and preparation of each sample, and the marker shape corresponds to the bioreactor. The percentages in parentheses represent the fraction of variance explained by that axis.

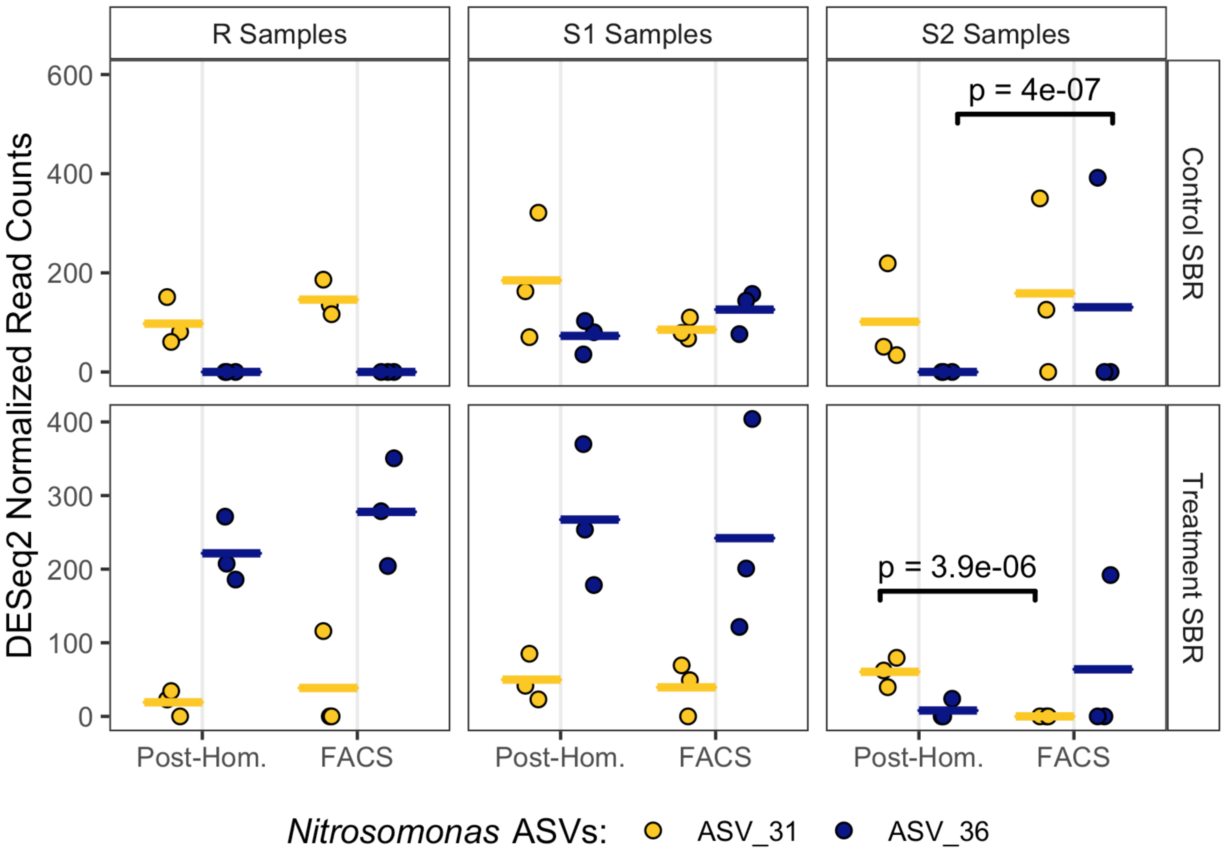

**Figure S12:** Normalized read counts of two predominant *Nitrosomonas* ASVs in triplicate BONCAT-FACS (i.e. BONCAT+) and corresponding Post-Homogenized (i.e. Post-Hom.) libraries generated from samples prepared from nitrifying microcosms seeded with mixed-liquor preceding return sludge treatment (R), return sludge after 15 min of side-stream treatment (S1), and return sludge after 24 h of side-stream treatment (S2) from the treatment SBR (bottom) and control SBR (top). Points indicate triplicate normalized read counts per ASV, and the horizontal bars of the same colors represent the sample mean per ASV. The reads were normalized with DESeq2 v.1.24.0 based on total read-counts per sample. A black bracket between an ASV within two samples represents a significant difference in the mean read abundance, determined with DESeq2 at an adjusted significance level of $p < 0.01$. These were the only two *Nitrosomonas* ASVs which showed differential abundance between BONCAT+ and corresponding Post-Hom. sample.

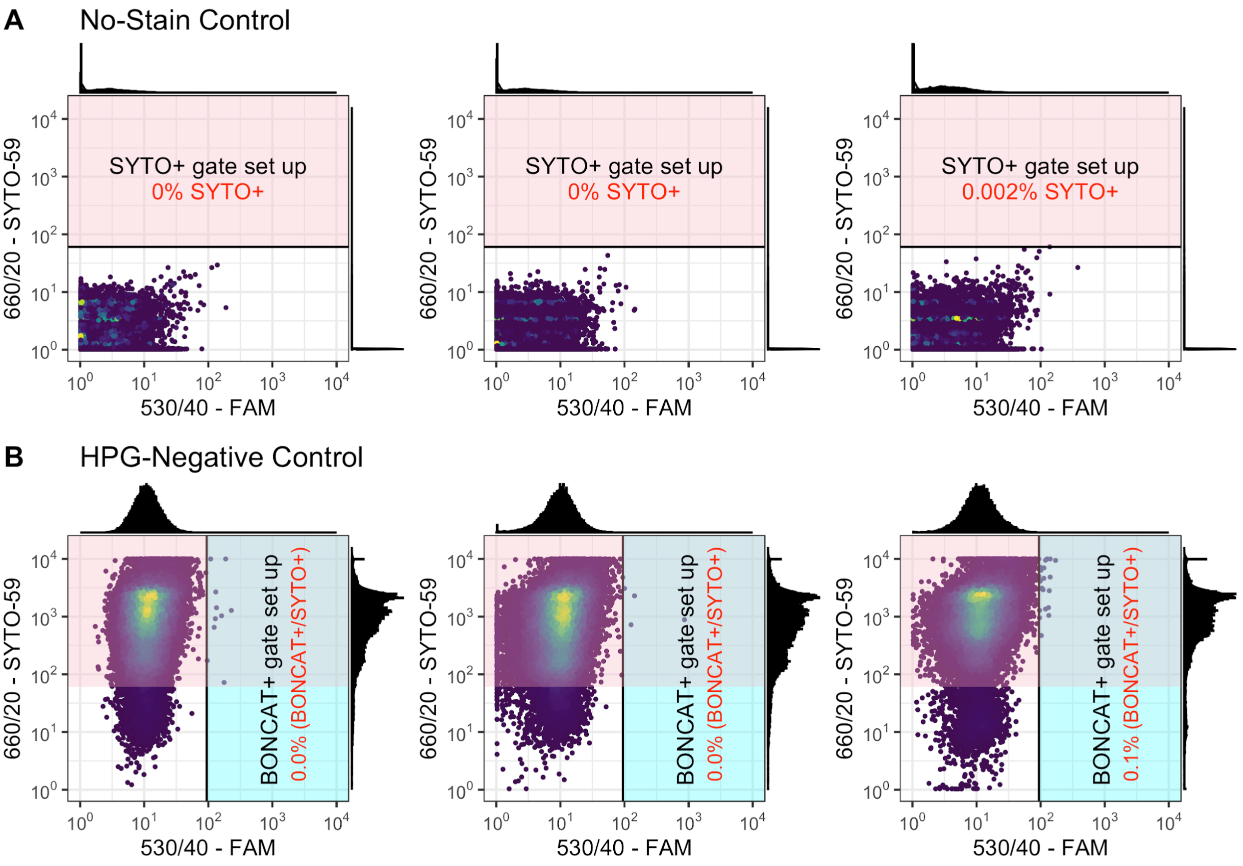

**Figure S13:** BONCAT-FACS gating setup using treatment SBR microcosm samples that were (A) not stained to determine background fluorescence passing a 660/20 filter; and (B) not incubated with HPG, but were exposed to click-chemistry with FAM picolyl azide dye and counter stained with SYTO59 to determine background fluorescence passing a 530/40 filter. The threshold for the SYTO+ and BONCAT+ regions were selected so that the mean false positive rate was below 0.5%.

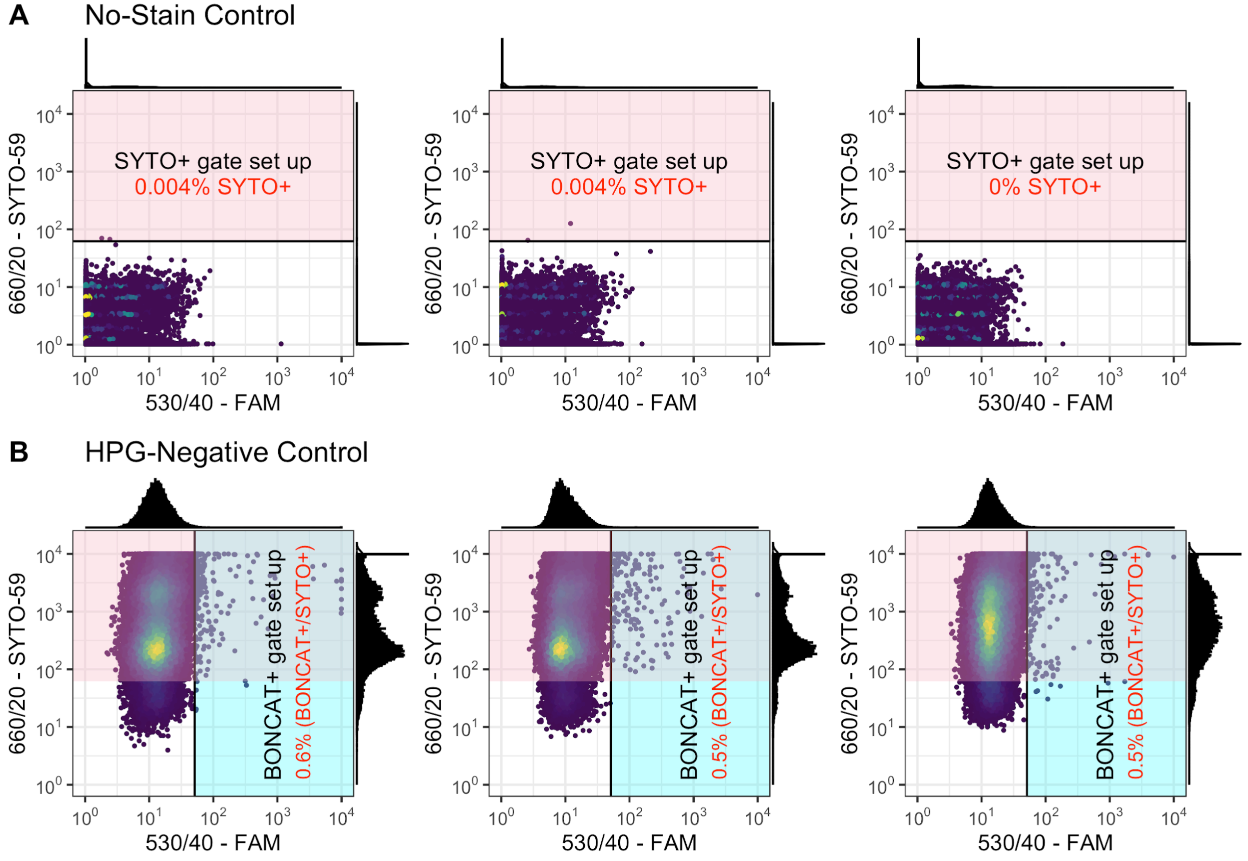

**Figure S14:** BONCAT-FACS gating setup using control SBR microcosm samples that were (A) not stained to determine background fluorescence passing a 660/20 filter; and (B) not incubated with HPG, but were exposed to click-chemistry with FAM picolyl azide dye and counter stained with SYTO59 to determine background fluorescence passing a 530/40 filter. The threshold for the SYTO+ and BONCAT+ regions were selected so that the mean false positive rate was below 0.5%.

**Table S1:** Summary of sequence read counts through amplicon filtering, denoising, merging, and chimera filtering.

| Sample ID | Reactor | Day | Input reads (F and R) | Filtered Reads (F and R) | Denoised F Reads | Denoised R Reads | Merged Reads | Non-chimeric Merged Reads |
|---|---|---|---|---|---|---|---|---|
| A1 | Treatment | -30 | 158970 | 146780 | 144148 | 144219 | 130317 | 94478 |
| B1 | Treatment | -30 | 37289 | 31734 | 31160 | 30964 | 28114 | 25731 |
| C1 | Treatment | -30 | 30772 | 27550 | 26785 | 26726 | 23852 | 21000 |
| D1 | Control | -30 | 30790 | 26971 | 26396 | 26253 | 24189 | 21624 |
| E1 | Control | -30 | 30919 | 27544 | 26853 | 26707 | 23962 | 21425 |
| F1 | Control | -30 | 31964 | 28396 | 27702 | 27652 | 25242 | 22848 |
| A3 | Treatment | -18 | 22201 | 20474 | 19793 | 19609 | 17576 | 15427 |
| B3 | Treatment | -18 | 17775 | 16369 | 15864 | 15703 | 14358 | 12964 |
| C3 | Treatment | -18 | 43335 | 39533 | 38769 | 38693 | 35757 | 32193 |
| D3 | Control | -18 | 29630 | 27223 | 26657 | 26490 | 24647 | 22802 |
| E3 | Control | -18 | 20393 | 18765 | 18261 | 18187 | 16803 | 15567 |
| F3 | Control | -18 | 14280 | 13202 | 12856 | 12819 | 11889 | 11368 |
| G1 | Treatment | 0 | 26019 | 19696 | 19203 | 19125 | 17595 | 16709 |
| H1 | Treatment | 0 | 33494 | 23238 | 22808 | 22664 | 21124 | 20268 |
| I1 | Treatment | 0 | 31909 | 24202 | 23695 | 23623 | 22228 | 20838 |
| J1 | Control | 0 | 29443 | 25079 | 24632 | 24466 | 22884 | 21733 |
| K1 | Control | 0 | 36891 | 27743 | 27430 | 27357 | 26185 | 25742 |
| L1 | Control | 0 | 19828 | 15570 | 15169 | 15155 | 14366 | 13864 |
| G3 | Treatment | 9 | 23618 | 21232 | 20736 | 20602 | 19129 | 17071 |
| H3 | Treatment | 9 | 33432 | 30730 | 30023 | 29813 | 27749 | 24557 |
| I3 | Treatment | 9 | 36653 | 33406 | 32691 | 32509 | 30274 | 27043 |

| | | | | | | | | |
|---|---|---|---|---|---|---|---|---|
| J3 | Control | 9 | 33094 | 30568 | 29753 | 29662 | 27227 | 23779 |
| K3 | Control | 9 | 28708 | 26672 | 25800 | 25833 | 23415 | 20497 |
| L3 | Control | 9 | 17751 | 16129 | 15470 | 15405 | 13688 | 12242 |
| A2 | Control | 31 | 58882 | 52946 | 52086 | 51966 | 48427 | 42035 |
| B2 | Control | 31 | 30516 | 25963 | 25390 | 25262 | 23471 | 21116 |
| M1 | Treatment | 31 | 36009 | 29146 | 28438 | 28344 | 25849 | 22393 |
| N1 | Treatment | 31 | 46647 | 40332 | 39294 | 39092 | 35323 | 30888 |
| O1 | Treatment | 31 | 49972 | 37597 | 36830 | 36693 | 33561 | 29125 |
| P1 | Control | 31 | 41192 | 36857 | 36062 | 35818 | 32862 | 27676 |
| A4 | Control | 46 | 38935 | 35194 | 34497 | 34259 | 31896 | 29163 |
| B4 | Control | 46 | 24131 | 15482 | 15066 | 14987 | 13897 | 12754 |
| M3 | Treatment | 46 | 24594 | 21508 | 20933 | 20707 | 18631 | 17446 |
| N3 | Treatment | 46 | 29108 | 26300 | 25824 | 25611 | 23699 | 22981 |
| O3 | Treatment | 46 | 38906 | 32246 | 31628 | 31365 | 28860 | 27324 |
| P3 | Control | 46 | 38920 | 35271 | 34571 | 34502 | 32070 | 29208 |
| C2 | Treatment | 61 | 24917 | 22698 | 21913 | 21769 | 18521 | 16196 |
| D2 | Treatment | 61 | 41147 | 33247 | 32527 | 32155 | 28557 | 26058 |
| E2 | Treatment | 61 | 40171 | 33306 | 32508 | 32349 | 28961 | 26690 |
| F2 | Control | 61 | 38625 | 28652 | 28285 | 28282 | 27158 | 26579 |
| G2 | Control | 61 | 51126 | 39764 | 39259 | 39063 | 36860 | 34181 |
| H2 | Control | 61 | 31152 | 15694 | 15327 | 15264 | 14492 | 14049 |
| C4 | Treatment | 75 | 88472 | 74762 | 73493 | 73271 | 67110 | 57452 |
| D4 | Treatment | 75 | 59213 | 53948 | 52990 | 52772 | 48153 | 43037 |

| | | | | | | | | |
|---|---|---|---|---|---|---|---|---|
| E4 | Treatment | 75 | 44888 | 39888 | 39138 | 38994 | 35853 | 33372 |
| F4 | Control | 75 | 35879 | 32832 | 32076 | 31924 | 29262 | 25730 |
| G4 | Control | 75 | 50487 | 45833 | 45090 | 45009 | 42047 | 38204 |
| H4 | Control | 75 | 40168 | 36511 | 35714 | 35684 | 32994 | 30227 |
| I2 | Treatment | 84 | 38826 | 28327 | 27682 | 27324 | 24273 | 22479 |
| J2 | Treatment | 84 | 28753 | 25008 | 24627 | 24436 | 22761 | 22207 |
| K2 | Treatment | 84 | 29252 | 25307 | 24583 | 24445 | 21182 | 19315 |
| L2 | Control | 84 | 24389 | 22166 | 21658 | 21557 | 19964 | 18058 |
| M2 | Control | 84 | 37453 | 32845 | 32326 | 32242 | 30582 | 29256 |
| N2 | Control | 84 | 26365 | 23983 | 23405 | 23206 | 21462 | 19697 |
| PN_1 | Treatment | 94 | 39294 | 30219 | 29070 | 29282 | 24892 | 15460 |
| PN_10 | Treatment | 94 | 98283 | 28177 | 27443 | 27445 | 24427 | 19329 |
| PN_11 | Treatment | 94 | 171219 | 4272 | 4045 | 4054 | 3493 | 3266 |
| PN_12 | Treatment | 94 | 120793 | 2792 | 2617 | 2602 | 2204 | 1784 |
| PN_13 | Treatment | 94 | 48022 | 33359 | 32286 | 32514 | 27040 | 17552 |
| PN_14 | Treatment | 94 | 45583 | 38811 | 37541 | 37727 | 31500 | 18490 |
| PN_15 | Treatment | 94 | 54551 | 33960 | 33105 | 32860 | 28133 | 21331 |
| PN_16 | Treatment | 94 | 53727 | 43267 | 42149 | 42401 | 37044 | 27284 |
| PN_17 | Treatment | 94 | 55579 | 46398 | 44960 | 44959 | 37820 | 25257 |
| PN_18 | Treatment | 94 | 60379 | 51320 | 49363 | 49692 | 41179 | 25902 |
| PN_19 | Treatment | 94 | 42281 | 35398 | 33620 | 33881 | 26637 | 16778 |
| PN_2 | Treatment | 94 | 77193 | 52459 | 50653 | 50884 | 42464 | 25129 |
| PN_20 | Treatment | 94 | 51432 | 35473 | 34818 | 34795 | 31268 | 26196 |

| | | | | | | | | |
|---|---|---|---|---|---|---|---|---|
| PN_21 | Treatment | 94 | 45181 | 33121 | 31930 | 32124 | 28017 | 18587 |
| PN_22 | Treatment | 94 | 55804 | 30123 | 29305 | 29338 | 26168 | 20837 |
| PN_23 | Treatment | 94 | 53350 | 45589 | 43883 | 44032 | 37357 | 24304 |
| PN_24 | Treatment | 94 | 29399 | 18536 | 17891 | 17985 | 15807 | 10917 |
| PN_25 | Treatment | 94 | 39641 | 31444 | 30847 | 30943 | 28102 | 23718 |
| PN_26 | Treatment | 94 | 54911 | 43159 | 42701 | 42694 | 39400 | 34466 |
| PN_27 | Treatment | 94 | 38564 | 32634 | 32051 | 32208 | 29573 | 24746 |
| PN_28 | Control | 94 | 39553 | 32552 | 31614 | 31744 | 28567 | 20187 |
| PN_29 | Control | 94 | 38319 | 29683 | 28852 | 28861 | 25891 | 17635 |
| PN_3 | Treatment | 94 | 54443 | 43258 | 41303 | 41908 | 33145 | 18015 |
| PN_30 | Control | 94 | 44397 | 36243 | 35261 | 35334 | 32096 | 22560 |
| PN_31 | Control | 94 | 40982 | 33041 | 32344 | 32585 | 30634 | 24832 |
| PN_32 | Control | 94 | 55275 | 44109 | 43144 | 42964 | 39261 | 29301 |
| PN_33 | Control | 94 | 45872 | 37984 | 37044 | 37294 | 34112 | 24852 |
| PN_34 | Control | 94 | 60718 | 32401 | 31906 | 31935 | 29840 | 26164 |
| PN_35 | Control | 94 | 71410 | 23690 | 23356 | 23321 | 22049 | 19307 |
| PN_36 | Control | 94 | 54342 | 4660 | 4391 | 4419 | 3829 | 3019 |
| PN_37 | Control | 94 | 50529 | 23683 | 23162 | 23126 | 21399 | 15362 |
| PN_38 | Control | 94 | 41924 | 32470 | 31563 | 31636 | 28665 | 20722 |
| PN_39 | Control | 94 | 45108 | 36495 | 35407 | 35451 | 31769 | 20264 |
| PN_4 | Treatment | 94 | 58814 | 50402 | 48770 | 49258 | 40474 | 22752 |
| PN_40 | Control | 94 | 86353 | 21121 | 20699 | 20472 | 18236 | 15548 |
| PN_41 | Control | 94 | 49851 | 38467 | 37215 | 37328 | 32586 | 23207 |

| | | | | | | | | |
|---|---|---|---|---|---|---|---|---|
| PN_42 | Control | 94 | 48507 | 39665 | 37828 | 38081 | 30724 | 18884 |
| PN_43 | Control | 94 | 40037 | 19425 | 19007 | 19060 | 17765 | 17437 |
| PN_44 | Control | 94 | 43166 | 37137 | 35391 | 35443 | 27744 | 16178 |
| PN_45 | Control | 94 | 45567 | 39198 | 37561 | 37745 | 31484 | 20106 |
| PN_46 | Control | 94 | 47815 | 39995 | 38746 | 38833 | 34598 | 22838 |
| PN_47 | Control | 94 | 55508 | 46974 | 45670 | 45732 | 41256 | 27398 |
| PN_48 | Control | 94 | 34451 | 28958 | 28161 | 28140 | 25480 | 17160 |
| PN_49 | Control | 94 | 57547 | 30062 | 29779 | 29740 | 28590 | 22207 |
| PN_5 | Treatment | 94 | 54988 | 36000 | 35234 | 35297 | 31352 | 22417 |
| PN_50 | Control | 94 | 66628 | 54408 | 53534 | 53688 | 49754 | 34614 |
| PN_51 | Control | 94 | 71856 | 29427 | 29066 | 28990 | 27327 | 23045 |
| PN_52 | Control | 94 | 56228 | 46180 | 45625 | 45725 | 43492 | 37648 |
| PN_53 | Control | 94 | 92789 | 24274 | 23797 | 23682 | 21574 | 19252 |
| PN_54 | Control | 94 | 51906 | 42149 | 40791 | 40681 | 34750 | 28061 |
| PN_55 | Treatment | 94 | 50130 | 41489 | 39784 | 39963 | 32428 | 17994 |
| PN_56 | Treatment | 94 | 68025 | 41706 | 40121 | 40519 | 33978 | 19507 |
| PN_57 | Treatment | 94 | 52995 | 37179 | 35864 | 35953 | 30007 | 17978 |
| PN_58 | Control | 94 | 71612 | 27296 | 26863 | 26811 | 25333 | 21383 |
| PN_59 | Control | 94 | 57208 | 26692 | 26250 | 26183 | 24631 | 21139 |
| PN_6 | Treatment | 94 | 58840 | 48307 | 47184 | 47194 | 40744 | 25431 |
| PN_60 | Control | 94 | 43926 | 29513 | 28852 | 28850 | 26605 | 18411 |
| PN_61 | Treatment | 94 | 60549 | 35224 | 34185 | 34395 | 29788 | 20274 |
| PN_62 | Treatment | 94 | 58423 | 42289 | 40399 | 40776 | 32882 | 17924 |

| | | | | | | | | |
|---|---|---|---|---|---|---|---|---|
| PN_63 | Treatment | 94 | 43627 | 30003 | 28770 | 28901 | 24295 | 14672 |
| PN_64 | Control | 94 | 55518 | 44069 | 42868 | 43091 | 38961 | 25005 |
| PN_65 | Control | 94 | 49792 | 40178 | 39049 | 39134 | 34442 | 22500 |
| PN_66 | Control | 94 | 54593 | 34704 | 33603 | 33703 | 29791 | 19671 |
| PN_67 | Treatment | 94 | 29278 | 24985 | 24205 | 24073 | 20761 | 17618 |
| PN_67_R | Treatment | 94 | 71295 | 60236 | 56510 | 57422 | 43863 | 25918 |
| PN_68 | Treatment | 94 | 34203 | 29627 | 28746 | 28612 | 24714 | 21187 |
| PN_68_R | Treatment | 94 | 55305 | 48423 | 45821 | 46276 | 35303 | 19451 |
| PN_69 | Treatment | 94 | 46014 | 39502 | 38516 | 38321 | 33902 | 29547 |
| PN_69_R | Treatment | 94 | 45467 | 39926 | 37549 | 37943 | 28911 | 17452 |
| PN_7 | Treatment | 94 | 42319 | 34412 | 33624 | 33734 | 30074 | 24347 |
| PN_70 | Control | 94 | 42722 | 36868 | 36159 | 35983 | 33393 | 29278 |
| PN_70_R | Control | 94 | 52901 | 45935 | 44208 | 44356 | 38957 | 28412 |
| PN_71 | Control | 94 | 27423 | 23478 | 22940 | 22811 | 21179 | 18867 |
| PN_71_R | Control | 94 | 36021 | 31277 | 29800 | 30064 | 25798 | 16892 |
| PN_72 | Control | 94 | 13602 | 11733 | 11399 | 11304 | 10341 | 9481 |
| PN_72_R | Control | 94 | 45556 | 39831 | 38034 | 38296 | 32883 | 18991 |
| PN_8 | Treatment | 94 | 46133 | 33318 | 32430 | 32397 | 28298 | 23038 |
| PN_9 | Treatment | 94 | 50655 | 26516 | 26120 | 26120 | 24627 | 22359 |

**Table S2:** Summary of analytical methods used for monitoring bioreactors

| Analyte | Method | Sample Pretreatment | Instrument |
|---|---|---|---|
| TSS | Standard Method 2540 D. Total Suspended Solids Dried at 103–105°C | Filtered onto glass microfiber filters (0.45-µm pore size, Whatman, Pittsburgh, USA), and dried in oven for 12 h | Fisher Scientific Isotemp 737F Oven for drying; Ohaus Adventurer AR2140 Analytical Balance (Ohaus, Parsippany, NJ, USA) for weighing |
| VSS | Standard Method 2540 E. Fixed and Volatile Solids Ignited at 550°C | Ignite residue from TSS measurement in furnace for 30 min | Thermolyne 30400 Furnace (Thermo Fisher Scientific) for burning; Ohaus Adventurer AR2140 Analytical Balance (Ohaus, Parsippany, NJ, USA) for weighting |
| Ammonia ($NH_3$-N) | Standard Method 4500-$NH_3$ H. Flow Injection Analysis | Centrifuged at 5000 rpm for 5 min, filtered through glass filters (0.2-µm pore size, Fisherbrand™), and stored at 4 °C before measurement | Thermo Iec Multi Rf Centrifuge (Thermo IEC, USA); Lachat QuikChem 8000 Series (Lachat Instrument, Wisconsin) |
| Nitrite ($NO_2^-$-N) | Standard Method 4500-$NO_2^-$ I. Cadmium Reduction Flow Injection Method | Centrifuged at 5000 rpm for 5 min, filtered through glass filters (0.2-µm pore size, Fisherbrand™), and stored at 4 °C before measurement | Thermo Iec Multi Rf Centrifuge (Thermo IEC, USA); Lachat QuikChem 8000 Series (Lachat Instrument, Wisconsin) |
| Nitrate ($NO_3^-$-N) | Standard Method 4500-$NO_3^-$ I. Cadmium Reduction Flow Injection Method | Centrifuged at 5000 rpm for 5 min, filtered through glass filters (0.2-µm pore size, Fisherbrand™), and stored at 4 °C before measurement | Thermo Iec Multi Rf Centrifuge (Thermo IEC, USA); Lachat QuikChem 8000 Series (Lachat Instrument, Wisconsin) |
| Orthophosphate ($PO_4^{3-}$-P) | Standard Method 4500-P G. Flow Injection Analysis for Orthophosphate | Centrifuged at 5000 rpm for 5 min, filtered through glass filters (0.2-µm pore size, Fisherbrand™), and stored at 4 °C before measurement | Thermo Iec Multi Rf Centrifuge (Thermo IEC, USA); Lachat QuikChem 8000 Series (Lachat Instrument, Wisconsin) |

| | | | |
|---|---|---|---|
| sCOD | N/A | Centrifuged at 5000 rpm for 5 min, filtered through glass filters (0.2-µm pore size, Fisherbrand™), acidified to pH 2 by $H_2SO_4$, and stored at 4 °C before measurement | Thermo Iec Multi Rf Centrifuge (Thermo IEC, USA); HP 6890 Series GC system (Hewlett Packard (Agilent)) |
| pH | N/A | Take less than 10 mL of mixed liquor from SBRs into beakers | Beckman 40 pH Meter (Beckman Coulter, Massachusetts, USA) |
| DO | N/A | N/A | HQ30D Portable DO Meter (Hach, Colorado, USA) |

**Text S1: Supplemental Methods**

*Micro and macro ingredients of synthetic wastewater:*

The synthetic wastewater contained macro elements (162.4 mg/L of $MgCl_2$, 128.65 mg/L of $CaCl_2 \cdot 2H_2O$, 79.1 mg/L of KCl), trace elements (1.03 mg/L of $FeSO_4 \cdot 7H_2O$, 0.32 mg/L of $ZnSO_4 \cdot 7H_2O$, 0.055 mg/L of $CuSO_4 \cdot 5H_2O$, 0.0562 mg/L of $CoCl_2 \cdot 6H_2O$, 0.032 mg/L of $Na_2MoO_4 \cdot 2H_2O$, 0.05 mg/L of $H_3BO_3$, 0.05 mg/L of KI, 0.022 mg/L of $NiCl \cdot 6H_2O$, 0.14 mg/L of $Al_2(SO_4)_3 \cdot 18H_2O$, 0.283 mg/L of $MnCl_2 \cdot 4H_2O$, 6.87 mg/L of EDTA), phosphate (51.3 mg/L of $Na_2HPO_4 \cdot 7H_2O$, 16.7 mg/L of $KH_2PO_4$), alkalinity (298.3 mg/L of $NaHCO_3$), and yeast (20 mg/L).

*Analytical methods for reactor monitoring:*

Please see Table S2 for a summary of analytical methods used for bioreactor monitoring.

*Daily effluent monitoring experiments to determine composite nitrite accumulation ratios:*

Monitoring experiments lasting 24 h were performed approximately every 10 days during the Treatment Phase to monitor daily nitrification performance over 8 of the 3 h SBR cycles. Mixed liquor samples were collected just after the addition of treated return sludge during the feeding period of the first cycle, and effluent samples were obtained during the settling period of the 1$^{st}$, 2$^{nd}$, 3$^{rd}$, 4$^{th}$, and 8$^{th}$ cycles thereafter. The nitrite accumulation ratio (NAR) of each cycle was calculated based on Equation 1, where $S_{NO2}$ and $S_{NO3}$ are the effluent nitrite and nitrate concentrations (mg-N/L), respectively.

**Eqn. 1**

$$NAR = S_{NO2} \div ( S_{NO2} + S_{NO3})$$

*Procedure and analysis of preliminary BONCAT validation microcosms:*

Three preliminary BONCAT microcosms were conducted prior to the experimental microcosms to validate the sensitivity of BONCAT in labelling only active cells that had incorporated HPG. The BONCAT validation microcosms included: (1) a HPG-negative control; (2) a pre-incubation fixed control; and (3) a post-incubation fixed control. All validation microcosms were seeded with mixed liquor from the control SBR and conducted exactly as the experimental microcosms except with the following modifications. For the pre-incubation fixed control, mixed liquor was first fixed with 3% paraformaldehyde for 1 h at RT and washed three times in 1x PBS prior to incubation with HPG. For the post-incubation fixed control, unfixed mixed liquor was first incubated with HPG before undergoing the same paraformaldehyde fixation and PBS washing steps. The HPG-negative control microcosm was conducted exactly as those in the experimental microcosms. Samples from each validation microcosm were washed three times in 1x PBS, resuspended in 50% ethanol in 1x PBS (vol/vol) and stored at -20°C prior to undergoing identical sample preparation and FACS analysis as the experimental microcosm samples.

*Homogenization procedure for BONCAT microcosm samples:*

The enzymatic homogenization procedure described below was optimized to enable the extraction of single microbial cells from complex activated sludge floc with minimal cell destruction. An 800 µL volume from each microcosm sample was transferred to a 15 mL Falcon tube, washed with 4mL of 1x PBS and resuspended into 2 mL of a filter-sterilized enzymatic homogenization solution consisting of 1 mg/mL protease (Protease from Streptomyces griseus, Type XIV, >3.5 units/mg, MilliporeSigma, USA) and 0.25 mg/mL α-amylase (Alpha-amylase from Bacillus sp., ~380 U/mg, MilliporeSigma) in 1x PBS. Samples were incubated in a temperaturecontrolled chamber at 37 °C with 200 rpm orbital shaking for 1.5 h. Samples were taken from the chamber and vortexed for 20 s at maximum speed every 0.5 h (three times total) throughout the incubation period. After the incubation period, 8 mL of a cold (4 °C) filter-sterilized solution of 10 mM EDTA in 1x PBS was added to each sample before incubating samples on ice for 15 min. Samples were then sonicated in an ice-filled sonicating water bath for three 1 min bursts with a 30 s rest period between each burst. Samples were stored in an ice-bath prior to filter-immobilization and click-chemistry labeling (conducted within 1 h).

*Click-chemistry labeling of homogenized BONCAT microcosm samples:*

Filter-immobilized click-chemistry was performed closely following the procedure outlined by Couradeau et al.(6) and in general adherence to the procedure outlined by Hatzenpichler and Orphan(7). Multiple aliquots of the click-solution were prepared by mixing the dye premix with the reaction buffer(6, 7). Aliquots of the dye-premix were prepared fresh by mixing 1.25 uL of 20 mM copper sulfate ($CuSO_4$, 100 uM final concentration), 2.50 μL of 50 mM tris-hydroxypropyltriazolylmethylamine (THPTA, 500 uM final concentration) and 1.67 μL of 1.5 mM FAM picolyl azide dye (10uM final concentration; Click Chemistry Tools, USA) and allowing the mixture to react for 3 min at RT in the dark. Aliquots of the reaction buffer were prepared by adding 12.5 μL of freshly prepared 100 mM sodium ascorbate in 1x PBS (5mM final concentration) and 12.5 μL of freshly prepared 100 mM aminoguanidine HCl in 1x PBS (5 mM final concentration) to 221 μL of 1x PBS. The dye-premix was added to the reaction buffer and the solution mixed gently by inversion to create the 'click-solution'.

Using a sterile syringe filtration apparatus, each homogenized microcosm sample (800 μL original volume) was immobilized onto a 0.22 um GTTP Isopore Membrane filter (MilliporeSigma, USA) and washed with 20 mL of filter-sterilized 1x PBS by passage through the filtration apparatus.

Filters were transferred onto glass slides and stored at RT in a light-shielded environment while the click-chemistry reaction mixture (click-solution) was prepared. Each filter-immobilized sample was covered with 80 µL of click-solution followed by a glass coverslip, and all samples were incubated in the dark for 30 min. Filter-immobilized samples were then washed three times in a series of 20 mL baths of 1x PBS for 5 min each to remove residual FAM dye. The washed filters were transferred into 15 mL Falcon tubes containing 2 mL of a filter-sterilized modified TE buffer (10mM Tris-Base, 5mM EDTA, pH 8) and three 2.3 mm zirconium/silica beads (11079125z, BioSpec, USA), and vortexed at 1800 rpm for 3.5 min to detach the cells from the filters. Each sample was filtered through a 30 µm Nylon Mesh filter (Nylon Net Filter, 30.0 um pore size, Millipore) using a sterile syringe filtration apparatus prior to SYTO59 counterstaining and FACS analysis (FACS conducted within 3 h of sample preparation). The No-stain control samples were prepared from HPG-negative microcosm samples using the same procedure but were omitted from click-chemistry labeling (no click-solution added) and SYTO59 counterstaining. Epifluorescence imaging was conducted to validate the efficacy of the sample homogenization and click-chemistry labeling procedures, and to provide visual confirmation of HPG-dependent BONCAT labeling (data not shown).

*Gating procedure for fluorescence activated cell sorting (FACS) of BONCAT-positive cells:*

Prepared (pre-sort) microcosm samples from each SBR were sorted in separate trials, with gates set independently in each trial to account for differences in biomass characteristics between the two SBRs. Initial gating was conducted using a gate drawn based on forward scatter and side scatter, followed by a gate drawn based on side scatter and trigger pulse width, to exclude large particles and cell aggregates  Sorting gates were established to simultaneously target SYTO59-positive (SYTO+) cells that were also BONCAT-positive (BONCAT+), as determined based on their FAM picolyl azide BONCAT dye fluorescence. A first gate was drawn to target STYO+ cells from background particles by analyzing the background SYTO59 fluorescence of cells in triplicate No-stain control samples (no click-chemistry, no SYTO59 counterstaining) and setting the gate such that < 0.5% false positives were included (Figures S2 and S3). A second gate was drawn to target BONCAT+ cells from non-HPG labelled cells by analyzing the background FAM fluorescence in triplicate HPG-negative control samples that underwent click-chemistry labeling and SYTO59 counterstaining, and setting the gate such that < 0.5% false positives were included (Figures S2 and S3). The sorting gate was then drawn as the area of overlap between the SYTO+ and BONCAT+ gates. The triplicate pre-sort samples from each HPG-positive microcosm were sorted based on their SYTO59 and FAM fluorescence using this predetermined sorting gate, where the fraction of BONCAT+ cells in each sample was calculated as the fraction of SYTO+ cells residing in the sorting gate.

*Low-biomass input DNA extraction procedure for BONCAT microcosm samples:*

The pre-homogenized, pre-sort, BONCAT-FACS samples prepared from each BONCAT microcosm (Section 2.3) were all extracted using the PrepGEM Bacteria kit (ZyGEM, New Zealand) following a procedure that was optimized for the extraction of the low biomass input BONCAT-FACS samples (i.e. low-biomass input procedure). The low-biomass input extraction procedure is outlined as follows. Samples were pulled from -80 °C storage and thawed on ice at 4 °C overnight. To each sample, 900 uL of ice-cold (-18 °C) 100% ethanol and 5.4 uL of 5mg/mL linear acrylamide (ThermoFisher Scientific, USA) were added before mixing samples by vortexing and incubating samples at -18 °C for 2 h to precipitate any DNA that may have been released by cell lysis during sample storage. The samples were then centrifuged (20,000 x g, 4 °C, 40 min) and the supernatant was removed through careful pipetting. Sample tubes were then opened and incubated in a heat block for 10 min at 37 °C to remove any residual supernatant through evaporation. A bulk volume of extraction mix was prepared following the prepGEM Bacteria kit instructions, and 10ul of the extraction mix was added to each sample before mixing samples briefly by vortexing. Extractions were then conducted by incubating samples in a heat block for 15 min at 37°C, 10 min at 75°C, and 2min at 92°C. The extracted samples were stored at 4°C prior to 16S rRNA gene amplification and sequencing.

*Estimating active biomass on VSS basis:*

Steady-state mass-balance equations for complete-mix AS with sludge recycle were used to predict the fraction of reactor VSS that was active biomass ($X_{bio}$) (8):

$$X_{bio} = X_H + X_{AOB} + X_{NOB}$$

$$X_{bio} = (SRT/HRT)[(Y_H)(COD_r)/(1 + b_H SRT) + (Y_{AOB})(NH_{4,ox})/(1 + b_{AOB} SRT) + (Y_{NOB})(NH_{4,ox})(1 - NAR)/(1 + b_{NOB} SRT)]$$

where:

*SRT* = solids retention time (10 d)
*HRT* = hydraulic retention time (0.5 d)
$Y_H$= heterotrophic yield (0.4 g VSS/g COD) (8)
$b_H$= heterotrophic decay constant (0.12 $d^{-1}$) (8)
$COD_r$ = COD removed (100 mg/L)
$Y_{AOB}$ = AOB yield (0.15 g VSS/g N) (8)
$b_{AOB}$= AOB decay constant (0.17 $d^{-1}$) (8)
$NH_{4,ox}$= amount of ammonium oxidized (~85% of influent (8) = 43 mg N/L flow-weighted average in FA phase)
$Y_{NOB}$ = NOB yield (0.05 g VSS/g N) (8)
$b_{NOB}$= NOB decay constant (0.17 $d^{-1}$) (8)
$NAR$= nitrite accumulation ratio ($NO_2^-/(NO_3^-+NO_2^-)$), used to infer fraction of un-oxidized $NO_2$-N (11% in Treatment SBR, 1% in control SBR)

With the above parameters, $X_{bio}$is estimated to be 426 mg VSS/L and 427 in the Treatment and control SBRs, respectively, which account for 77% and 67% of the total reactor VSS, respectively (Figure S2). This analysis assumes the decay constants were similar in two reactors, which may not have been the case.

*Bioinformatics script for denoising 16S rRNA gene amplicon data with DADA2:*

The following markdown was run in the R environment (v.3.6.1). The code chunks in grey background were run for this analysis.

```
###1. Load libraries

library(dada2); packageVersion("dada2")

library(tidyverse);

In this study, we had 2 different MiSeq runs on amplicons targeting the V4-V5 region of the 16S rRNA gene. Because error rates may be different for each run, we will do denoising, and merging on a run-by-run basis. But, to ensure the same amplicon size, we will filter all reads together.

Is this R-markdown being run for the first time, or would you like to override previously saved data?

###2. Quality filter reads

ovr <- "FALSE" #change to TRUE if you want to override data

Set path directory to where raw_seqs folder is stored
#Import file names
path <- "./16S_iTags" # CHANGE ME to the directory containing the analysis files.
path_fs <- paste0(path,"/raw_seqs")

Read in sample metadata:
sampledata_pn <- read.csv(file=paste0(path,"/metadata.csv"), header=TRUE, row.names=1, stringsAsFactors = FALSE) %>% #read in sample metadata from plate submission to sequencing facility
  dplyr::filter(Reactor == "R1" | Reactor =="R2") %>% #filter metadata to just select PN samples
  mutate(Sample = paste0(Plate_Column, Plate_Row)) %>% #use plate row and column to make sample names like the .fastq files
    mutate(Sample = str_replace_all(Sample, "-", "_"))
tail(sampledata_pn)
Read in file names
fns <- list.files(path_fs)
fns
fastqs <- fns[grepl("001.fastq$", fns)]
fastqs <- sort(fastqs) # Sort ensures forward/reverse reads are in same order
fnFs <- fastqs[grepl("_R1", fastqs)] # Just the forward read files
fnRs <- fastqs[grepl("_R2", fastqs)] # Just the reverse read files
```

Fully specify the path for the fnFs and fnRs

```
fnFs <- file.path(path_fs, fnFs)
fnFs
fnRs <- file.path(path_fs, fnRs)
```

### 3. Filter and trim reads

Visualizing the quality profiles along the sequencing reads

```
plotQualityProfile(fnFs[[1]])
plotQualityProfile(fnRs[[1]])
```

Make directory and filenames for the filtered fastqs:

```
filt_path <- file.path(path, "filtered")
if(!file_test("-d", filt_path)) dir.create(filt_path)
if(ovr) filtFs <- file.path(filt_path, paste0(sample.names, "_F_filt.fastq.gz")) #create placeholder files for filtered reads
if(ovr) filtRs <- file.path(filt_path, paste0(sample.names, "_R_filt.fastq.gz"))
names(filtFs) <- sample.names
names(filtRs) <- sample.names
```

Filter sequences by trimming first 10bp, truncating F reads at 275bp and R reads at 225bp

```
if(ovr) out <- filterAndTrim(fnFs, filtFs, fnRs, filtRs, trimLeft=c(15, 15), truncLen=c(275,225),maxN=0, maxEE=c(2,2), truncQ=2, rm.phix=TRUE,compress=TRUE, multithread=TRUE)
# On Windows set multithread=FALSE
if(ovr) saveRDS(out, file=paste0(path,"/out.RDS")) #save R object in case we need to re-run later
if(file_test("-f", paste0(path,"/out.RDS"))) out <- readRDS(file=paste0(path,"/out.RDS"))
```

Learn error rates of forward reads for run 1

```
if(ovr) errF1 <- learnErrors(filtFs1, multithread=TRUE)
if(ovr) saveRDS(errF1, file=paste0(path,"/errF1.RDS")) #save R object in case we need to re-run later
if(file_test("-f", paste0(path,"/errF1.RDS"))) errF1 <- readRDS(file=paste0(path,"/errF1.RDS"))
```

Learn error rates of reverse reads for run 1

```
if(ovr) errR1 <- learnErrors(filtRs1, multithread=TRUE)
if(ovr) saveRDS(errR1, file=paste0(path,"/errR1.RDS")) #save R object in case we need to re-run later
if(file_test("-f", paste0(path,"/errR1.RDS"))) errR1 <- readRDS(file=paste0(path,"/errR1.RDS"))
```

Learn error rates of forward reads for run 2

```
if(ovr) errF2 <- learnErrors(filtFs2, multithread=TRUE)
if(ovr) saveRDS(errF2, file=paste0(path,"/errF2.RDS")) #save R object in case we need to re-run later
if(file_test("-f", paste0(path,"/errF2.RDS"))) errF2 <- readRDS(file=paste0(path,"/errF2.RDS"))
```

Learn error rates of reverse reads for run 2

```
if(ovr) errR2 <- learnErrors(filtRs2, multithread=TRUE)
if(ovr) saveRDS(errR2, file=paste0(path,"/errR2.RDS")) #save R object in case we need to re-run later
```

```
if(file_test("-f", paste0(path,"/errR2.RDS"))) errR2 <-
readRDS(file=paste0(path,"/errR2.RDS"))
```

Plot errors

```
plotErrors(errF1, nominalQ=TRUE)
plotErrors(errF2, nominalQ=TRUE)
plotErrors(errR1, nominalQ=TRUE)
plotErrors(errR2, nominalQ=TRUE)
```

Sample inference using above model fit, run1

```
#Forward reads
if(ovr) dadaFs1 <- dada(filtFs1, err=errF1, multithread=TRUE)
if(ovr) saveRDS(dadaFs1, file=paste0(path,"/dadaFs1.RDS")) #save R
object in case we need to re-run later
if(file_test("-f", paste0(path,"/dadaFs1.RDS"))) dadaFs1 <-
readRDS(file=paste0(path,"/dadaFs1.RDS"))

#Reverse reads
if(ovr) dadaRs1 <- dada(filtRs1, err=errR1, multithread=TRUE)
if(ovr) saveRDS(dadaRs1, file=paste0(path,"/dadaRs1.RDS")) #save R
object in case we need to re-run later
if(file_test("-f", paste0(path,"/dadaRs1.RDS"))) dadaRs1 <-
readRDS(file=paste0(path,"/dadaRs1.RDS"))
```

Sample inference using above model fit, run2

```
#Forward reads
if(ovr) dadaFs2 <- dada(filtFs2, err=errF2, multithread=TRUE)
if(ovr) saveRDS(dadaFs2, file=paste0(path,"/dadaFs2.RDS")) #save R
object in case we need to re-run later
if(file_test("-f", paste0(path,"/dadaFs2.RDS"))) dadaFs2 <-
readRDS(file=paste0(path,"/dadaFs2.RDS"))

#Reverse reads
if(ovr) dadaRs2 <- dada(filtRs2, err=errR2, multithread=TRUE)
if(ovr) saveRDS(dadaRs2, file=paste0(path,"/dadaRs2.RDS")) #save R
object in case we need to re-run later
if(file_test("-f", paste0(path,"/dadaRs2.RDS"))) dadaRs2 <-
readRDS(file=paste0(path,"/dadaRs2.RDS"))
```

### 4. Merge paired reads

Check that samples are in right order

```
if(!identical(names(filtFs1), names(filtRs1))) stop("Forward and
reverse files do not match.")
if(!identical(names(filtFs2), names(filtRs2))) stop("Forward and
reverse files do not match.")
```

Run 1

```
if(ovr) mergers1 <- mergePairs(dadaFs1, filtFs1, dadaRs1, filtRs1,
verbose=TRUE)
```

Run 2

```
if(ovr) mergers2 <- mergePairs(dadaFs2, filtFs2, dadaRs2, filtRs2,
verbose=TRUE)
```

Make sequence tables:

Run 1

```
if(ovr) seqtab1 <- makeSequenceTable(mergers1)
```

```
if(ovr) saveRDS(seqtab1, file=paste0(path,"/seqtab1.RDS")) #save R
object in case we need to re-run later
if(file_test("-f", paste0(path,"/seqtab1.RDS"))) seqtab1 <-
readRDS(file=paste0(path,"/seqtab1.RDS"))
dim(seqtab1)
Run 2
if(ovr) seqtab2 <- makeSequenceTable(mergers2)
if(ovr) saveRDS(seqtab2, file=paste0(path,"/seqtab2.RDS")) #save R
object in case we need to re-run later
if(file_test("-f", paste0(path,"/seqtab2.RDS"))) seqtab2 <-
readRDS(file=paste0(path,"/seqtab2.RDS"))
dim(seqtab2)

### 5. Remove chimeras
if(ovr) seqtab <- mergeSequenceTables(seqtab1, seqtab2)
if(ovr) saveRDS(seqtab, file=paste0(path,"/seqtab.RDS")) #save R object
in case we need to re-run later
if(file_test("-f", paste0(path,"/seqtab.RDS"))) seqtab <-
readRDS(file=paste0(path,"/seqtab.RDS"))

if(ovr) seqtab.nochim <- removeBimeraDenovo(seqtab, method="consensus",
multithread=TRUE, verbose=TRUE)
if(ovr) saveRDS(seqtab.nochim, file=paste0(path,"/seqtab_nochim.rds"))
if(file_test("-f", paste0(path,"/seqtab_nochim.rds"))) seqtab.nochim <-
readRDS(file=paste0(path,"/seqtab_nochim.rds"))
```
